# Studying ecosystems with DNA metabarcoding: lessons from aquatic biomonitoring

**DOI:** 10.1101/578591

**Authors:** Alex Bush, Zacchaeus Compson, Wendy Monk, Teresita M. Porter, Royce Steeves, Erik Emilson, Nellie Gagne, Mehrdad Hajibabaei, Mélanie Roy, Donald J. Baird

## Abstract

An ongoing challenge for ecological studies has been the collection of data with high precision and accuracy at a sufficient scale to detect effects relevant to management of critical global change processes. A major hurdle for many workflows has been the time-consuming and challenging process of sorting and identification of organisms, but the rapid development of DNA metabarcoding as a biodiversity observation tool provides a potential solution. As high-throughput sequencing becomes more rapid and cost-effective, a ‘big data’ revolution is anticipated, based on higher and more accurate taxonomic resolution, more efficient detection, and greater sample processing capacity. These advances have the potential to amplify the power of ecological studies to detect change and diagnose its cause, through a methodology termed ‘Biomonitoring 2.0’.

Despite its promise, the unfamiliar terminology and pace of development in high-throughput sequencing technologies has contributed to a growing concern that an unproven technology is supplanting tried and tested approaches, lowering trust among potential users, and reducing uptake by ecologists and environmental management practitioners. While it is reasonable to exercise caution, we argue that any criticism of new methods must also acknowledge the shortcomings and lower capacity of current observation methods. Broader understanding of the statistical properties of metabarcoding data will help ecologists to design, test and review evidence for new hypotheses.

We highlight the uncertainties and challenges underlying DNA metabarcoding and traditional methods for compositional analysis, focusing on issues of taxonomic resolution, sample similarity, taxon misidentification, sample contamination, and taxon abundance. Using the example of freshwater benthic ecosystems, one of the most widely-applied non-microbial applications of DNA metabarcoding to date, we explore the ability of this new technology to improve the quality and utility of ecological data, recognising that the issues raised have widespread applicability across all ecosystem types.

## Introduction

Biodiversity loss and the risks it poses to ecosystem functions and services remain a major societal concern (Cardinale et al. 2012), but due to a lack of consistently-observed data, there is no consensus regarding the speed or severity of this decline (Vellend et al. 2013; Newbold et al. 2015). There are very few ecosystems in which we can quantify the magnitude of degradation, nor can we discriminate among multiple stressors, both key goals for environmental monitoring programs (Bonada et al. 2006). The power to detect change in ecological communities has been hampered by sampling costs predominantly associated with human labour and travel. As a result, ecosystem monitoring programs must manage a trade-off between the scope of a study, including the phylogenetic breadth of taxon coverage and the resolution to which taxa are described, and its spatial and temporal coverage (e.g. tropical forests Gardner et al. 2008; marine sediments Musco et al. 2009). A history of such trade-offs has led to entrenched practices relying on observation of a narrow range of taxa, which aim to provide a surrogate for the full biodiversity complement, yet whose taxonomic, spatial or temporal relationships are largely undefined (Lindenmayer & Likens 2011). The troubling reality is that management decisions are informed by very limited and potentially biased information, generated by approaches that no longer reflect our understanding of how ecosystems and species interact (Woodward, Gray & Baird 2013).

Fortunately, technological advances offer the opportunity to generate high-quality biodiversity data in a consistent manner, radically expanding the scope of ecosystem monitoring (e.g. Turner 2014; Bush et al. 2017). One of the most promising of these is the technique of DNA metabarcoding, which supports the massively-parallelised taxonomic identification of organism assemblages within a biological sample. The application of this method in ecosystem monitoring, termed “Biomonitoring 2.0” (Baird & Hajibabaei 2012) uses this approach to support the generation of higher level ecological knowledge that supports advances in our understanding of metacommunity and food-web theory (Bohan et al. 2017). When fully realised, DNA metabarcoding will provide a universal platform to identify any, and potentially all, phylogenetic groups occurring within an ecosystem, including many taxa currently not identifiable by expert taxonomists (e.g. streams: Sweeney et al. 2011; rainforest: Brehm et al. 2016; marine zooplankton: Zhang et al. 2018). As DNA sequencing capacity continues to increase, there is a growing interest from ecological researchers and environmental managers for guidance in how to apply these new tools, and to provide clear evidence of their value relative to existing microscopy-based methods. However, it is important to emphasise that comparisons between traditional morphological identifications and DNA sequences are far from straightforward. For example, while metabarcoding can observe the occurrence of DNA sequences within a specified environmental matrix (e.g. soil sample), it does not currently discriminate between intact, living organisms and their presence as parts, ingested, or extraneous tissue. While some may see this as a challenge to be overcome, to retrofit a new method to an old system of observation, we view this as an opportunity to expand our universe of interest and gain new insight into ecosystem structure and function (Bohan et al. 2017). Using data from our own and other studies, we explore the uncertainties surrounding both traditional and DNA-based observation approaches. Our examples are drawn largely from recent research on river ecosystems, a research area with a long history and strong linkages with regulatory application for assessing the state of the environment (Friberg et al. 2011; Leese et al. 2018).

Aquatic researchers have long recognised the challenges of taxonomic identification and resulting limitations it imposes on the scale and scope of observational, experimental and monitoring studies (Jones 2008). Freshwater monitoring programs rely upon a subset of taxa, primarily aquatic macroinvertebrates, fish, or algae, with little consistency across environmental agencies or regions (Friberg et al. 2011), and sparse spatial and temporal coverage and limited taxonomic resolution (e.g. Orlofske & Baird 2013) ultimately constrains outcomes to ‘pass/fail’ (impacted/non-impacted; Clarke et al. 2006; Strachan & Reynoldson 2014), with causes of degradation inferred rather than supported by direct evidence. After decades of research, our ability to disentangle the influence of even the most basic drivers that impact the state of freshwater ecosystems is still limited (Woodward, Gray & Baird 2013).

## Our unit and universe of observation

The science of aquatic biomonitoring is based on the principle that site-level observations of biological assemblage structure integrate responses to prevailing environmental conditions over space and time, reducing the intensity of sampling required to detect stressor-related changes in the environment, and providing an immediate signal of “ecosystem health” (Friberg et al. 2011). However, consistently observing more than a narrow range of taxa within an ecological community has proved costly and impractical, with accuracy of identification often unrecorded or difficult to quantify, and varying across taxa. The observation universe is further constrained by sampling method (e.g. mesh-size of collection nets), rather than common phylogenetic or ecological characteristics, with further downgrading or exclusion of groups that are difficult to identify (e.g. Vlek, Šporka & Krno 2006). Even with the best taxonomic expertise available, it is practically impossible to identify all specimens to species-level, since many early life-stages lack necessary diagnostic features (Orlofske & Baird 2013). Species are subsequently aggregated at higher taxonomic ranks, obscuring species-level responses, constraining our knowledge of whether species’ environmental preferences are conserved or variable (Macher et al. 2016; Beermann et al. 2018). In our view, the level of observation provided by direct morphological identification of biological specimens in a sample is highly variable (typically referred to as “lowest taxonomic level”), disconnected from ecological theory, and contains an unknown yet potentially significant degree of bias (Jones 2008).

Ecological field studies inevitably face budgetary constraints, and DNA metabarcoding offers the potential to reduce many of the costs involved in routine morphological identification (Ji et al. 2013). While single-specimen DNA barcoding uses short genetic sequences to identify individual taxa, often at the species-level, metabarcoding supports simultaneous identification of entire assemblages of organism via high-throughput sequencing (Taberlet et al. 2012; Yu et al. 2012). Metabarcoding has now been applied in a wide range of aquatic ecosystems (e.g. rivers: Hajibabaei et al. 2011; wetlands: Gibson et al. 2015; lakes: Bista et al. 2017) and used to describe community composition in a wide variety of taxa (e.g. worms: Vivien et al. 2015; insects: Emilson et al. 2017; diatoms: Vasselon et al. 2017).

When combined with appropriate bioinformatics tools, DNA-based identification can generate lists of taxa that are typically far richer than those generated by morphological identification (Sweeney et al. 2011; Gibson et al. 2015). This is further enhanced by expanding DNA barcode reference libraries (e.g. Curry et al. 2018) and by machine-learning algorithms (Porter & Hajibabaei 2018c). This has the potential to remove a significant impediment in field ecological studies, which need no longer be constrained by available taxonomic expertise. This new observation paradigm supports a definable universe of observation based on the types of DNA barcodes sequenced (see also below).

## Defining the universe of observation with metabarcoding

While metabarcoding offers the potential to observe a greater diversity of freshwater taxa, the requirement to amplify extracted DNA to generate sufficient material for sequencing places limitations on simultaneous, universal taxonomic observation. The selection of primers used to amplify specific DNA sequence marker regions is crucial to any metabarcoding study, since they are necessarily tailored to the taxonomic groups under study (Hajibabaei et al. 2012; Gibson et al. 2014). In order to expand taxonomic coverage, it is necessary to employ a range of primers and marker sequences (see Fig.3 in Gibson et al. 2014). Considerable efforts have been made to develop and refine primers for different taxonomic groups or species, and primers with broad coverage for invertebrates have now been established (e.g. Hajibabaei et al. 2012; Elbrecht & Leese 2017). However, amplification bias due to variable affinity among sequence variants for amplification can distort the relationship between sample biomass and the number of sequence reads (Elbrecht & Leese 2015; Zhang et al. 2018). Metabarcoding can therefore support a taxonomically broad universe of observation, but outputs should be treated as occurrences and do not currently support reliable estimation of organism biomass or abundance.

Before discussing the parallels and differences between morphology-based monitoring and metabarcoding, two key issues must be highlighted: the distinction between bulk-community sampling and environmental DNA (eDNA), and the choice of primers. eDNA samples focus on a signal derived predominantly from traces of intracellular and extracellular DNA without attempting to isolate organisms (e.g. from water or soil; Cristescu & Hebert 2018), whereas bulk-community samples include eDNA, but target the collection of whole organisms. eDNA can be effective in detecting biological signal from the environment, but the significant spatial and temporal uncertainty of that signal clouds its application in observational studies. As a result, our examples of metabarcoding below focus entirely on observations derived from bulk-community samples that are otherwise identical to traditional monitoring surveys.

## Interpretation

The statistical power and precision of any ecological assessment based on sample assemblage composition depends upon how results are combined and scored, and how identification errors (i.e. false-presences and false-absences) can obscure the calibration of baseline composition, limiting our ability to detect deviations from this baseline and infer that change has occurred (e.g. Clarke et al. 2002; Clarke 2009). Although many sources of uncertainty affect our ability to infer regional and landscape-level trends from site-level observations, these are difficult to address with traditional approaches (Clarke 2009; Carstensen & Lindegarth 2016). To illustrate this problem, we focus on how five sources of error involved in describing freshwater biodiversity differ between morphological and metabarcoding workflows: a) taxonomic resolution, b) replicate similarity, c) taxonomic misidentification, d) contamination, and e) quantitative measures like abundance.

### Taxonomic resolution

Biomonitoring 2.0 (Baird & Hajibabaei, 2012) employs metabarcoding to overcome the taxonomic bottleneck of sample processing, removing a critical trade-off between sample taxonomic resolution and the number of samples that can be studied (Jones 2008). Moreover, sample metrics derived from higher taxonomic categories, such as family- or genus-level, make a tacit assumption that species within those higher categories share similar environmental responses, and possess similar ecological functions. However, when studies are able to differentiate taxa at the species level, this assumption is false (e.g. nutrient and sediment sensitivity; Macher et al. 2016; Beermann et al. 2018), and this can significantly influence study outcomes (Hawkins et al. 2000; Schmidt-Kloiber & Nijboer 2004; Sweeney et al. 2011).

Observing taxonomic assemblages at genus- or family-level masks turnover in composition, reducing our power to detect subtle changes among communities over space and time. As each species is less common than its parent taxonomic group, there will be fewer observations with which to establish reliable associations, and their inclusion could add noise to statistical models, echoing the long-running debate about the value of rare taxa in biomonitoring (Nijboer & Schmidt-Kloiber 2004). This “noise” is not only due to the stochastic occurrence of uncommon species, but also sampling error, which can be quantified before discarding data (Clarke 2009; Ficetola, Taberlet & Coissac 2016; Guillera-Arroita 2016). We should therefore be particularly cautious about concluding how taxonomic resolution affects the strength of statistical relationships (Arscott, Jackson & Kratzer 2006; Martin, Adamowicz & Cottenie 2016). Instead, our current challenge is understanding when these subtle changes, previously invisible to traditional monitoring, are related to natural environmental factors or anthropogenic disturbance.

One criticism of DNA metabarcoding is that high taxonomic resolution is not valuable if those taxa cannot be linked to a binomial taxonomic name, a limitation that emerges when barcode reference libraries are incomplete (Curry et al. 2018). However, many methods of ecological assessment evaluate community level characteristics such as alpha- and beta-diversity, that do not retain taxon identity, particularly at the species-level (Pawlowski et al. 2018). For this reason, interest in taxonomy-free approaches is increasing among those studying poorly-known assemblages whose morphological identification is challenging (e.g. meiofauna or diatoms: Vasselon et al. 2017). Moreover, new metrics could improve compatibility between biogeographically separated programs (Turak et al. 2017). Nonetheless, to tie DNA-based monitoring to historic surveys, and to assign ancillary information such as traits, it is still a requirement to assign taxonomic names to identified sequences (e.g. Compson et al. 2018). Based on the wealth of ecological information available that could complement DNA-based ecological studies, and the considerable body of legacy data generated by historical studies, including regulatory monitoring, increasing reference library coverage should be a priority for management agencies transitioning to DNA-based surveys.

### Replicate similarity

Depending on the scale of observation, species are rarely distributed randomly or uniformly in nature. For example, the distribution of macroinvertebrate taxa in streams is notoriously dynamic, as species adjust to changes in both abiotic (e.g. flow velocity, substratum size) and biotic (e.g. fish predation, mussel aggregation) factors (Downes, Lake & Schreiber 1993; Vaughn & Spooner 2006). Heterogeneity may also result from stochastic processes such as dispersal and colonization (Fonseca & Hart 2001), ephemeral resources (Lancaster & Downes 2014), or disturbance regimes at multiple scales (Effenberger et al. 2006). Indeed, heterogeneity is so pervasive that a shift towards greater homogeneity within aquatic communities could indicate human modification of the landscape (Petsch 2016). Given such heterogeneity, the challenge for ecological studies or biomonitoring is to detect a sufficient proportion of the community, whilst also minimising processing costs, so that further detections are unlikely to alter the interpretation of subsequent analyses. Counting all individuals in a sample can have value, but it is prohibitive for routine observational studies, and not cost-effective for biomonitoring purposes (e.g. Vlek, Šporka & Krno 2006). Most studies therefore employ subsampling (i.e. identifying a subset of individuals collected from the field) to reduce the time, effort and cost of processing macroinvertebrate samples. However, reducing the effort per sampling unit can significantly underestimate the richness per sample (Doberstein, Karr & Conquest 2000; Buss et al. 2014) and although subsampling is standardized by volume, weight, or number of individuals, it is often difficult to compare among survey methods and biomonitoring schemes (Buss et al. 2014). Although sensitivity to subsampling depends on the metric employed, subsampling can substantially increase the misclassification of site status (Clarke et al. 2006; Petkovska & Urbanič 2010) and exaggerate the perceived rarity of many taxa, whose exclusion from analyses may further bias interpretations of condition (Schmidt-Kloiber & Nijboer 2004).

Regardless of the sub-sampling approach, a single sample only recovers a subset of the community, particularly in heterogeneous environments (Fig. 1 & 2). As sampling effort increases, either by area or time, more taxa are recovered until the rate of new discoveries declines (Vlek, Šporka & Krno 2006). The rate of accumulation depends on taxon abundance distributions, their dispersion, and ease of collection, including the effects of environment on collection efficiency (Guillera-Arroita 2016). For example, a typical 3-minute kick-sample recovered only 50% of the macroinvertebrates species, and 60% of the families, found in total from six replicate samples (Furse et al. 1981). Other standardized protocols observe a similar degree of turnover among replicates (Fig. 1).

**Figure 1.**
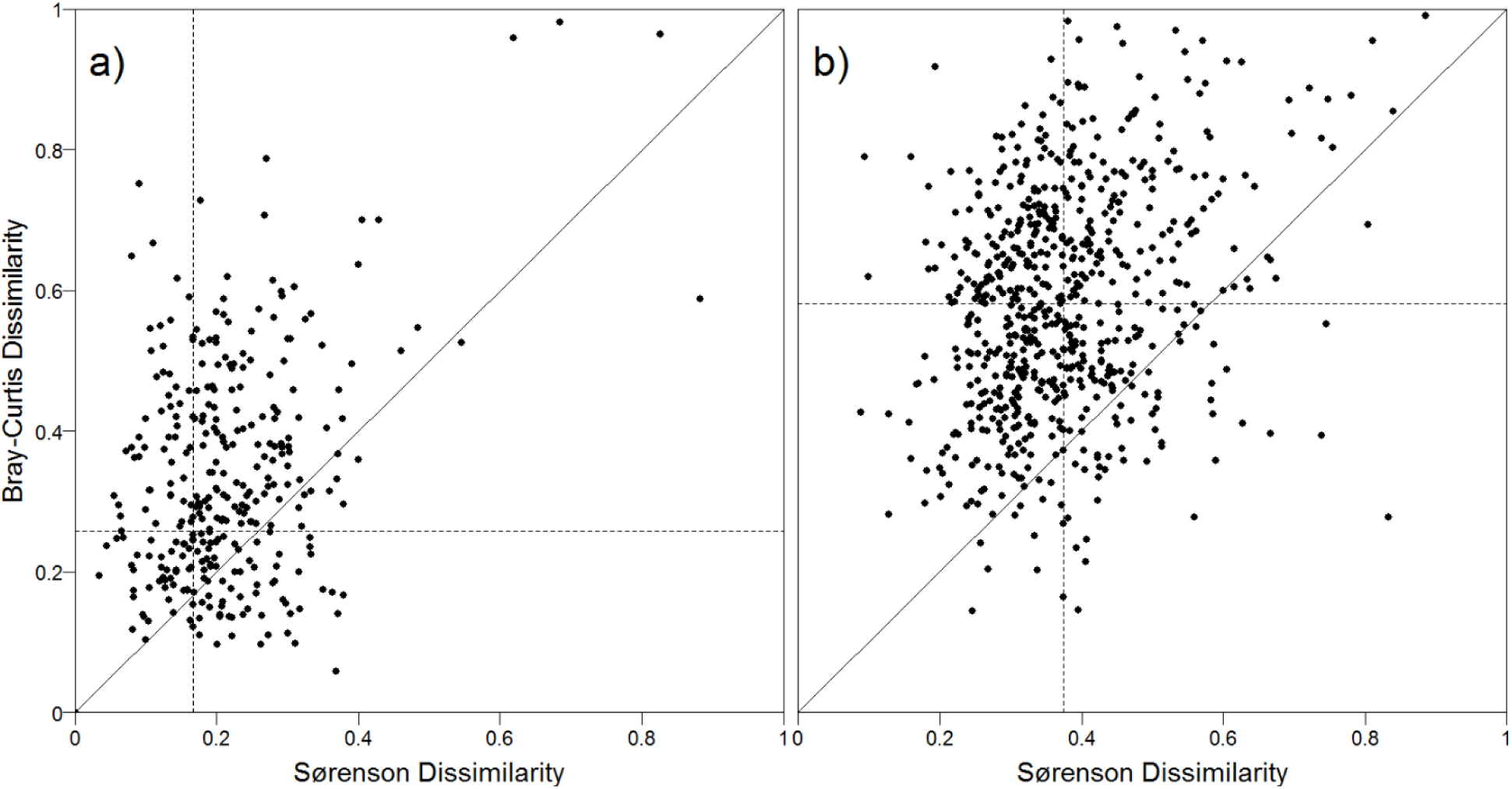
Dissimilarity between replicate samples based on presence/absence data (Sørensen), and count data (Bray-Curtis) of morphologically identified macroinvertebrate families from a) 417 CABIN (Canadian Aquatic Biomonitoring Network; ECCC 2018) surveys (total n=1656, mean richness=16+/-4.8), and b) 787 surveys from the STAR-AQEM dataset (total n=1673) from 14 European countries (mean richness=51 +/-18.4; (Furse et al. 2006; Schmidt-Kloiber et al. 2014).

Metabarcoding can, in principle, substantially reduce this sampling error, since the entire sample is processed (but see also limitations associated with primer selection discussed below). False absences can be further reduced by rarefying the number of taxa observed per read and by analysing technical replicates (i.e. multiple DNA aliquots from sample extracts). Although low-biomass, low abundance taxa may still be missed (Hajibabaei et al. 2012; Elbrecht, Peinert & Leese 2017), metabarcoding detects a higher proportion of the target assemblage compared to morphologically-identified samples (Fig. 2), thereby increasing the power of monitoring programs to detect change.

**Figure 2.**
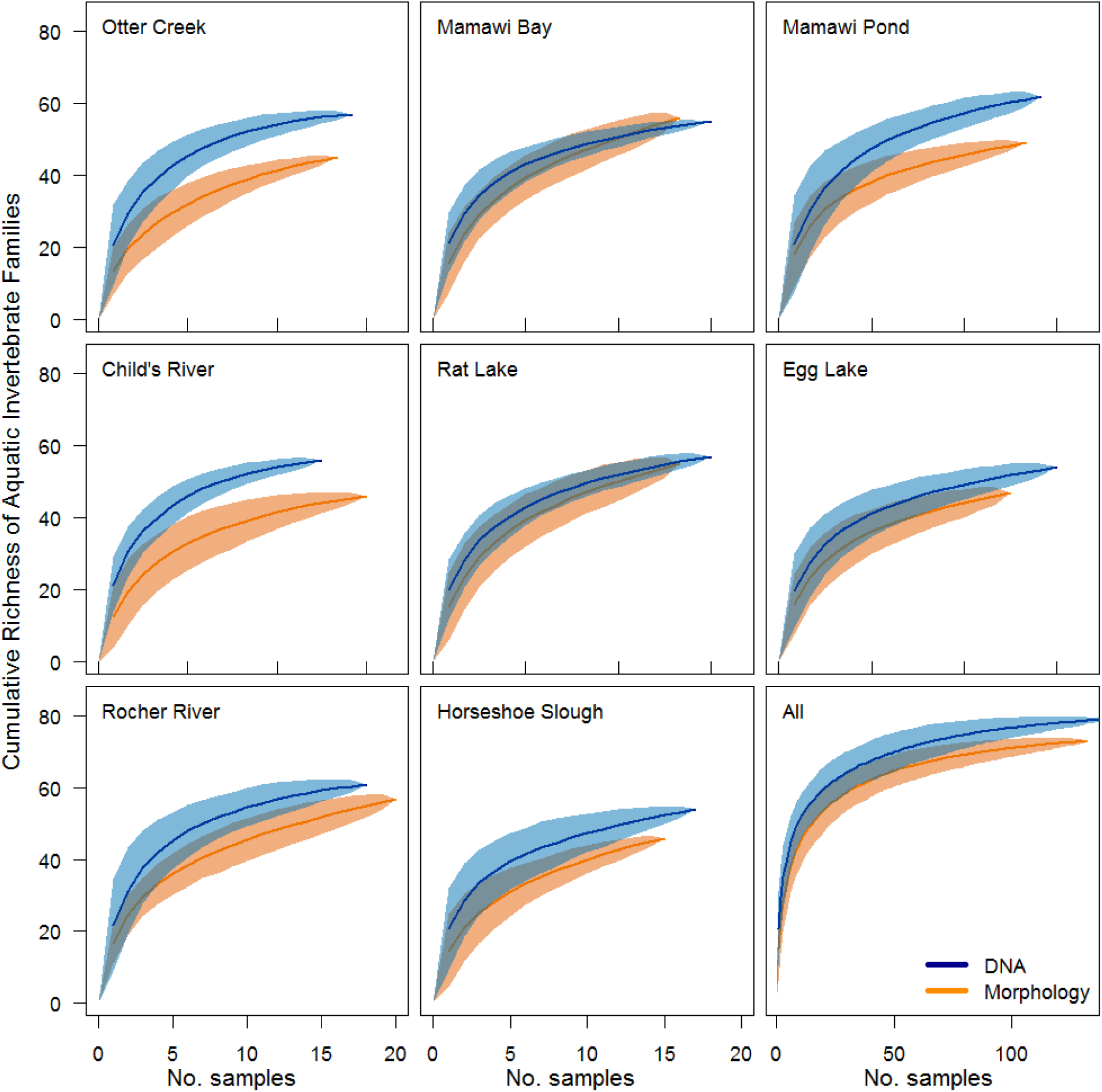
Accumulated richness (mean +/-95% confidence interval) of aquatic invertebrate families from 8 wetland sites in the Peace-Athabasca Delta, and for all samples combined (note different scale). Samples were collected between 2011 and 2016 (updated from surveys published in Gibson et al. 2015).

### Misidentification

Morphological identification of diverse taxonomic groups, such as invertebrates, is challenging, as demonstrated by a lack of reliable species-level data generated by routine biomonitoring programs. The probability of misidentifying an individual depends on the quality of the specimen (e.g. is the specimen partial or complete? Is it mature or immature?), the availability and completeness of identification keys, and the taxonomist’s experience. Though most biomonitoring programs now include a process for quality control and assessment to limit the likelihood of misidentification, false positives and negatives are still common. For example, early audits of the RIVPACS program showed that 8.3% of family occurrences were missed, and approximately one false presence was added in every four samples (Clarke 2009). Similarly, an audit of a range of European programs by Haase et al. (2006) found that after accounting for misidentifications and sorting errors, samples were on average 40% dissimilar to their initial composition. These errors compound the loss of taxa during sub-sampling, but remain difficult to predict.

A major advantage of metabarcoding over traditional morphological identification is the ability to generate accurate identifications in a consistent manner (Orlofske & Baird 2013; Jackson et al. 2014). That said, the accuracy of metabarcoding still depends on the taxonomic coverage and quality of reference DNA sequences used for taxonomic inference as well as the bioinformatics approaches employed (Porter & Hajibabaei 2018b). If organisms are misidentified at the time of sequence deposition, reference library sequences become associated with an incorrect taxonomic name. To minimise this challenge, the Barcode of Life Database (BOLD) stores information on voucher specimens, supporting linkage of sequences to material in curated reference specimen collections. Overall, database coverage for animals is expanding rapidly (Porter & Hajibabaei 2018b) and is already relatively high for freshwater invertebrates. For example, sequences exist for 95% of the genera observed in >1% of samples collected by the Canadian national biomonitoring program (Curry et al. 2018; see also Leese et al. 2018). The current BOLD reference library is better suited to identifying macroinvertebrate families routinely observed in Canada, reflecting the greater effort on DNA barcode library development in that country when compared to Australia and the UK (Figure 3). Consequently, at the time of writing, a routine Bayesian classifier (Porter & Hajibabaei 2018a) is expected to misidentify 4.4%, 6.1% and 7.7% of families within CABIN, RIVPACS and AUSRIVAS programs respectively. It cannot be overstated that this is a significant improvement on the documented ability of current best-available morphological identification, and is accompanied by an ability to drill down to species-level, which will only improve as DNA libraries become more complete. To further improve DNA-based identification by barcodes, agencies considering the transition to metabarcoding should support targeted specimen collection, and accelerate the digitisation of existing museum-collected material to improve geographic and taxonomic library coverage (Stokstad 2018).

**Figure 3.**
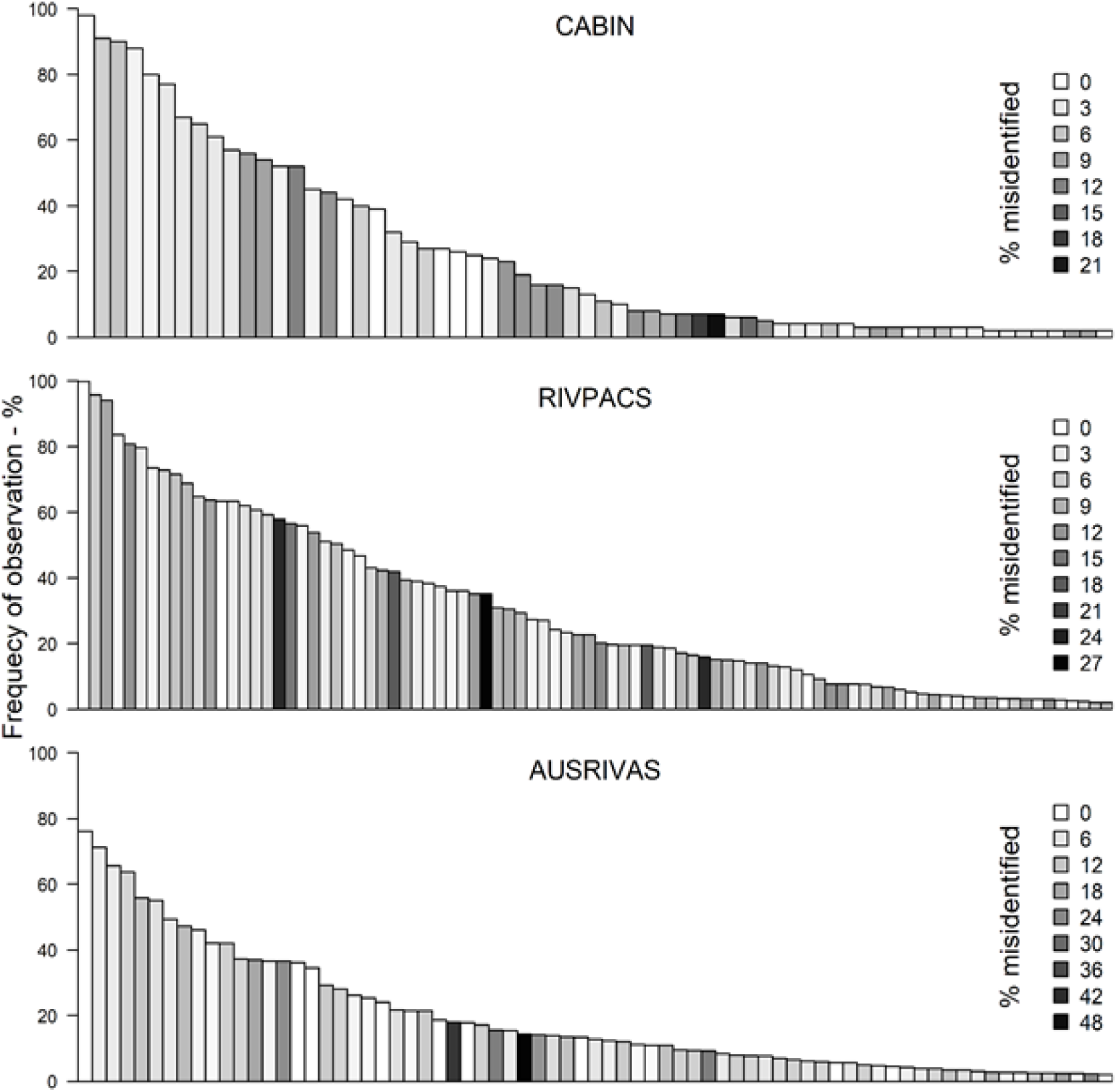
Families ordered by frequency of occurrence within three biomonitoring programs: the CABIN (*n* = 540), the UK River Invertebrate Prediction and Classification System (RIVPACS, n=2,504), and the Australian River Assessment System (AUSRIVAS *n*=1,516) from Victoria. Shading reflects the likelihood taxa could be misidentified using the CO1 RDP classifier v.3 (see Supplement 1 for further details).

### Contamination

The detection sensitivity of metabarcoding has raised concerns that the number of false positives will increase, particularly due to the adventitious introduction of DNA that did not originate from the sampled site. Existing ecological sampling protocols often recommend cleaning of equipment between surveys to reduce transfer of invasive species or pathogens, and a more rigorous version of this practice should be adopted as standard practice to reduce the possibility of cross-sample contamination with DNA. Quality control and assurance practices are particularly crucial in eDNA studies that amplify trace amounts of DNA; these studies often include various controls, such as samples from localities that are believed to lack the target taxa, extraction blanks, and equipment controls. A combination of replicate sampling and appropriate controls can then quantify the rate of false-positives and false-negatives before observations are confirmed (Ficetola et al. 2015). Thus, although it is difficult to eliminate the possibility of cross-contamination altogether, it is possible to greatly reduce its occurrence and precisely quantify the probability of errors to support study quality assurance and control.

### Quantitative measures of biodiversity

As stated above, DNA metabarcoding results do not currently produce a reliable signal of abundance or biomass (Elbrecht & Leese 2015). Nonetheless, it is equally misleading to suggest that current biomonitoring practices are themselves able to effectively detect differences in macroinvertebrate abundance without substantial effort. The difficulty of processing samples, coupled with species’ patchy distributions, means few studies can claim to have truly quantified patterns of abundance for multispecies invertebrate assemblages (e.g. Hawkins et al. 2000).

Obtaining a reliable estimate of taxon abundance or biomass can support studies of many key ecological processes, but for the specific purposes of detecting compositional change, abundance information is most useful when responses can indicate a shift in species dominance without a change in composition. This is particularly true in depauperate systems, if species are pooled at higher taxonomic levels, or rare taxa are discarded (Reynoldson et al. 1997). Nonetheless, differences in the composition of diverse assemblages are often sufficient to discriminate among sites, even at relatively coarse taxonomic resolution (Thorne, Williams & Cao 1999; Hawkins et al. 2000), thus the challenge has always been the reliable identification of those taxa. While count or relative abundance information may provide another axis for discrimination, its inherent variability exaggerates the dissimilarity among replicate samples (Fig. 1), rendering baseline conditions more variable, thus reducing statistical power to detect change. These limitations are well illustrated by studies that have replaced quantitative count data with qualitative categories or occurrence data (e.g. Wright et al. 1984; Armanini et al. 2013). These approaches have proved acceptable to practitioners precisely because count data provide little or no incremental improvement to detecting differences among sites. Moreover, approaches based on occurrence data illustrate a direct pathway to implement DNA metabarcoding in routine biomonitoring programs.

### Performance

The relative advantages of DNA metabarcoding over morphological methods are necessarily contingent on the nature and scope of the question being investigated. Bonada et al. (2006) reviewed the requirements of biomonitoring studies to detect the occurrence and intensity of anthropogenic impacts, and Dafforn et al. (2016) explored their applicability to answer questions over a range of spatial and temporal scales. As they are driven by regulatory needs, most monitoring programs focus on relatively simple outcomes (e.g. local deviation from baseline; categorical quality assessment), and thus can greatly benefit from increased precision and statistical power. Recent freshwater ecosystem studies have demonstrated that metabarcoding data can support detection of ecological change at a greater level of discrimination than traditional approaches (Gibson et al. 2015; Elbrecht et al. 2017; Emilson et al. 2017). Although regulators have thus far remained hesitant to transition to monitoring with metabarcoding, these early studies have highlighted a lack of precision and consistency in the application of existing morphological approaches, shortcomings of traditional morphological observation that too often are either ignored or unrecognized by current practitioners.

Our purpose in developing DNA metabarcoding as an observational tool has been to explore its ability to provide consistently-observed information to answer routine questions posed by managers (e.g. is biological composition at a site significantly different from expectations, and if so, is there evidence of impact?). Comparisons between metabarcoding and morphology-based methods have involved sorting and identification of a sample using existing taxonomic keys, followed by the reassembly of the sample for metabarcoding (Hajibabaei et al. 2012; but see Gibson et al. 2015). These approaches have demonstrated that DNA metabarcoding recovered ~90% of the taxa identified by morphology, and all false-absences were from taxa that represented <1% of individuals. Most recently, we have also evaluated the similarity of taxa recovered by metabarcoding using paired samples (Fig. 4; GRDI-Ecobiomics 2017). The average similarity of morphological and metabarcoded samples at family-level was 73%, within the range of variation expected for replicate samples (Fig. 1; Clarke et al. 2002). Of the families observed by both methods, DNA observed 79% of the observations made by morphology, whereas morphology only matched 61% of those made by DNA. Some families also appear to be consistently under-represented or absent from this DNA dataset (Fig.4a-b, bottom-left), most likely due to a combination of gaps in the reference library (aquatic mites and oligochaetes in particular) and primer bias (Gibson et al. 2014; Elbrecht et al. 2017). Beyond mere overlap, a better estimate of performance could be the likelihood each family was missed based on their detectability in replicate samples (Fig.4b). Both methods are likely to have missed many families at least once, but the mean and likelihood of multiple false absences was lower among metabarcoding samples than for samples identified by morphology (Supplement 2).

**Figure 4.**
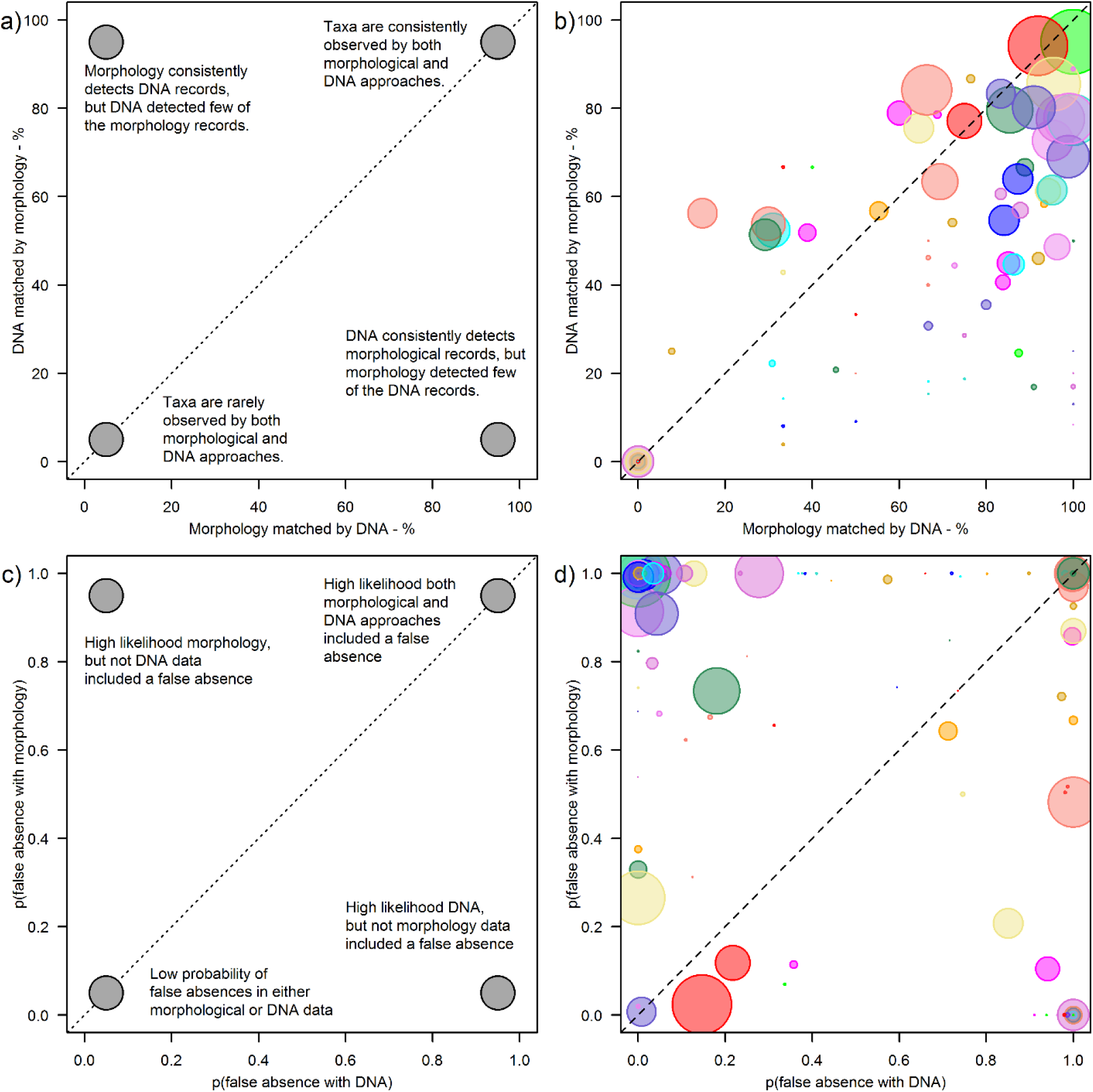
Comparison of macroinvertebrate families (n=114) observed in pairs of standard 3-minute river benthos kick samples (n=141 sites). Top row (a and b) shows the correspondence between observations of each taxonomic family using either morphological identification or DNA metabarcoding. Points are scaled relative to the number of morphological observations. Bottom row (c and d) shows the probability that each method included at least one false absence for each taxon (see Supplement 2 for code and raw data).

While primer bias remains an issue, the composition recovered by DNA metabarcoding is always likely to be a subset of all taxa in diverse systems. Nonetheless, metabarcoding provides a step-change in taxonomic coverage, in terms of the taxonomic breadth of taxa observed, improved taxonomic resolution, and fewer false negatives. Compared to traditional morphological methods, metabarcoding representing a major advance in how consistently we observe the taxonomic structure of ecological communities.

## Conclusions

Biomonitoring 2.0 (Baird & Hajibabaei 2012) envisaged the use of DNA metabarcoding to generate consistently-observed biodiversity data to detect environmental change efficiently and rapidly. This can be done with only minor modification of existing sample collection methods, ensuring backwards compatibility with legacy data. Higher taxonomic resolution, more efficient detection (Fig. 2), and the capacity to increase spatiotemporal coverage can all increase the statistical power to detect change and diagnose its cause (Bonada et al. 2006).

Study design and interpretation should acknowledge the sources of uncertainty in both morphological and metabarcoding approaches. Although abundance information is specified by existing programs (Leese et al. 2018), it is not necessary to achieve biomonitoring goals and many robust methods that use occurrence information already exist. Sources of uncertainty associated with metabarcoding can be quantified and minimised more easily than morphological approaches (e.g. Davis et al. 2018), and once standard operating procedures emerge, many tasks can be automated, further reducing the risk of handling errors and the costs of sequencing (Porter & Hajibabaei 2018c). A transition to large-scale observation by metabarcoding will take time as sequencing still requires specialized technicians and facilities. However, as demand grows, we anticipate organisations will outsource their DNA sample processing to specialist labs, equivalent to the current use of private consultants for taxonomic and chemical analyses. Currently, the cost of processing an invertebrate community sample (from DNA-extraction to sequencing) is approximately half the cost of morphological identification by taxonomists, but as we have stressed, the divergent properties of each approach make it misleading to base comparisons on costs alone.

We can only manage what we can measure, and at present the unknown magnitude and consequences of global biodiversity loss emphasize the value of metabarcoding as a technique to support improved ecological observation in all field studies of multispecies assemblages. Moving forward, we expect the increasing number of metabarcoding studies to further refine the uncertainties associated with observations, and the exchange of information should accelerate as research activities in this area grow, spearheading large-scale implementation of metabarcoding. Metabarcoding is also being used for increasingly novel applications, such as the study of trophic interactions (Bohan et al. 2017), meta-community theory (Miller, Svanbäck & Bohannan 2018), and ecosystem function relationships (Vamosi et al. 2017), and these applications could generate substantial added value to existing or future biomonitoring programs (Compson et al. 2018).

## Acknowledgements

We thank Guy Woodward, Richard Marchant, Astrid Schmidt-Kloiber and Daniel Hering for providing data from monitoring programs in the UK, Australia and EU. This work was supported by the Ontario Genomics Institute and Genome Canada, NSERC, Environment and Climate Change Canada program funds and the Canadian federal Genomics Research & Development Initiative.

