## Supplement 1 for "Studying ecosystems with DNA metabarcoding: lessons from aquatic biomonitoring"

### Supplementary Material S1 Likelihood macroinvertebrate families observed in freshwater biomonitoring can be identified using CO1 barcodes based on current reference library coverage.

*Alex Bush*

*18 December 2018*

#### RDP Classifier

Format data used by the RDP classifier to represent the taxonomic hierarchy and the bootstrap classification scores (*available on Github repository* **https://github.com/terrimporter/CO1Classifier**).

Below are some steps that were taken to format the files available with each version of the RDP classifier:

```
# RDP Taxonomy
rdp_hierarchy = read.delim("mytaxon.txt", sep="*")[,-4]
names(rdp_hierarchy) = c("tax_id","name_txt","parent_tax_id","Rank")

# Directory of output files from bootstrap testing (26 in all)
# Contains the bootstrap confidence intervals for the RDP Classifier V.3 based on 200bp fragments
bootdir   = "../TP_scripts/CO1_V3_bootstrap_results/loo_out"
bootfiles = list.files(bootdir, pattern = "\\.out$",  full.names=T)

for(i in 1:length(bootfiles)){
  v3dat = readLines(bootfiles[i])
  v3dat = v3dat[c((grep("\\*\\*misclassified sequences group by taxon",v3dat)+2):
                    (grep("\\*\\*ROC matrix",v3dat)-1))]
  v3dat = unlist(lapply(v3dat, function(x) strsplit(x, "\\\t")[[1]]))
  v3dat = data.frame("Rank"                          = v3dat[seq(1,length(v3dat),6)],
                     "Name"                          = v3dat[seq(2,length(v3dat),6)],
                     "Total_Seqs"                    = v3dat[seq(3,length(v3dat),6)],
                     "Tested_Seqs_nonsingleton"      = v3dat[seq(4,length(v3dat),6)],
                     "misclassified"                 = v3dat[seq(5,length(v3dat),6)],
                     "pct_misclassified"             = v3dat[seq(6,length(v3dat),6)] )
  v3dat$Total_Seqs               = as.numeric(as.character(v3dat$Total_Seqs))
  v3dat$Tested_Seqs_nonsingleton = as.numeric(as.character(v3dat$Tested_Seqs_nonsingleton))
  v3dat$misclassified            = as.numeric(as.character(v3dat$misclassified))
  v3dat$pct_misclassified        = as.numeric(as.character(v3dat$pct_misclassified))

  # Combine
  if(!exists("bootdat")){
    bootdat = v3dat
  } else {
    if(any(is.na(match(v3dat$Name,bootdat$Name)))){ stop("NA's") }
    # Add sequences tested and misclassified to existing table
    bootdat[,4:5] = bootdat[match(bootdat$Name,v3dat$Name),4:5] +
      v3dat[(v3dat$Name%in%bootdat$Name),4:5]
  }
```

```
}

bootdat$Name = as.character(bootdat$Name)
bootdat$tax_id = rdp_hierarchy$tax_id[match(bootdat$Name,rdp_hierarchy$name_txt)]
bootdat$parent_tax_id = rdp_hierarchy$parent_tax_id[match(bootdat$Name,
                                                          rdp_hierarchy$name_txt)]
# Recalculate the percentage misclassified
bootdat$pct_misclassified = bootdat$misclassified / (bootdat$Tested_Seqs_nonsingleton/100)

# I could switch taxa under "Plecoptera" for "Plecoptera_Insecta" in each dataset,
# but instead I will remove the entries in the classifier for the moth genus
# 'Plecoptera' (from SE Asia)
bootdat$Name = gsub("Plecoptera_Insecta","Plecoptera",bootdat$Name)
bootdat = bootdat[-c(grep("Plecoptera_",bootdat$Name)),]

# Save to file and reloaded below
save(bootdat,       file="Supplementary_Data_RDP_classification_success.RData")
save(rdp_hierarchy, file="Supplementary_Data_RDP_hierarchy.RData")
```

For the purposes this demo the tables are available as supplementary data:

```
load("Supplementary_Data_RDP_hierarchy.RData")
kable(head(rdp_hierarchy), booktabs=TRUE)
```

| tax_id | name_txt | parent_tax_id | Rank |
|---|---|---|---|
| 2 | Archaea | 1 | superkingdom |
| 3 | undef_Archaea | 2 | kingdom |
| 4 | Euryarchaeota | 3 | phylum |
| 5 | Halobacteria | 4 | class |
| 6 | Haloferacales | 5 | order |
| 7 | Halorubraceae | 6 | family |

```
load("Supplementary_Data_RDP_classification_success.RData")
kable(bootdat[3:8,], booktabs=TRUE)
```

| | Rank | Name | Total_Seqs | Tested_Seqs_nonsingleton | misclassified | pct_misclassified | tax_id | p |
|---|---|---|---|---|---|---|---|---|
| 3 | kingdom | Fungi | 1452 | 1311 | 273 | 20.82380 | 1540 | |
| 4 | phylum | Ascomycota | 1135 | 1049 | 227 | 21.63966 | 1541 | |
| 5 | class | Sordariomycetes | 224 | 199 | 34 | 17.08543 | 1942 | |
| 6 | order | Glomerellales | 43 | 40 | 9 | 22.50000 | 1953 | |
| 7 | family | Australiascaceae | 1 | 0 | 0 | NaN | 1954 | |
| 8 | genus | Monilochaetes | 1 | 0 | 0 | NaN | 1955 | |

---

#### CABIN

Load list of macroinvertebrate families observed in CABIN surveys, and the number and percentage of sites in which they were observed.

```r
load("Supplementary_Data_CABIN_Taxa_Frequency.RData")
```

Families that do not match the RDP taxonomy:

```r
kable(canX[is.na(match(canX$Family, bootdat$Name)),], booktabs=TRUE)
```

| | Family | No_sites | Perc_of_sites | Order | Class | Phylum |
|---|---|---|---|---|---|---|
| 1 | Aeolosomatidae | 1 | 0 | Haplotaxida | Clitellata | Annelida |
| 7 | Anisitsiellidae | 28 | 2 | Trombidiformes | Arachnida | Arthropoda |
| 13 | Aturidae | 44 | 4 | Trombidiformes | Arachnida | Arthropoda |
| 56 | Eylaidae | 4 | 0 | Trombidiformes | Arachnida | Arthropoda |
| 57 | Feltriidae | 7 | 1 | Trombidiformes | Arachnida | Arthropoda |
| 75 | Hydrodromidae | 2 | 0 | Trombidiformes | Arachnida | Arthropoda |
| 79 | Hydrozetidae | 41 | 4 | Sarcoptiformes | Arachnida | Arthropoda |
| 97 | Macrothricidae | 7 | 1 | Diplostraca | Branchiopoda | Arthropoda |
| 98 | Mideopsidae | 13 | 1 | Trombidiformes | Arachnida | Arthropoda |
| 101 | Naididae | 320 | 27 | Haplotaxida | Clitellata | Annelida |
| 106 | Oreoleptidae | 4 | 0 | Diptera | Insecta | Arthropoda |
| 107 | Oxidae | 16 | 1 | Trombidiformes | Arachnida | Arthropoda |
| 141 | Stygothrombiidae | 8 | 1 | Trombidiformes | Arachnida | Arthropoda |

Attach data from the RDP classifier on the percentage of times species in this Family were misidentified, and then reduce the table to those taxa for which we can estimate the rate of misclassification, or occur in fewer than 1% of sites.

```r
# Add column for the percent misclassified
canX$pct_misclassified = bootdat$pct_misclassified[match(canX$Family, bootdat$Name)]
# Remove any families that occurred in less than 1% of (reference) sites, or don't have a misclassifica
canX = canX[(canX$Perc_of_sites>0) & (!is.na(canX$pct_misclassified)),]
```

---

#### RIVPACS

Load list of macroinvertebrate families observed in RIVPACS surveys, and the number and percentage of sites in which they were observed.

```r
load("Supplementary_Data_RIVPACS_Taxa_Frequency.RData")
```

#### AUSRIVAS

Load list of macroinvertebrate families observed in AUSRIVAS surveys, and the number and percentage of sites in which they were observed.

```r
load("Supplementary_Data_AUSRIVAS_Taxa_Frequency.RData")
```

---

#### PLOTTING CO1 SEQUENCE COVERAGE OF MONITORING PROGRAMS

Combining the data from CABIN, RIVPACS AND AUSRIVAS we can now visualise how often Families commonly encountered by each monitoring program would be expected to be misidentified using DNA metabarcoding based on bootstrap-test statistics of the RDP classifier.

```
# Format a new joint table
bmidat = rbind(canX[,c("Family","Perc_of_sites","Order","pct_misclassified")],
               ukX[,c("Family","Perc_of_sites","Order","pct_misclassified")],
               ausX[,c("Family","Perc_of_sites","Order","pct_misclassified")])
bmidat$Program = c(rep("CABIN",nrow(canX)),rep("RIVPACS",nrow(ukX)),rep("AUSRIVAS",nrow(ausX)))
bmidat = bmidat[complete.cases(bmidat),]
# Remove Families is less than 2% of samples
bmidat = bmidat[-c(which(bmidat$Perc_of_sites<2)),]
# Sort data
bmidat = bmidat[order(bmidat$Family, bmidat$Order, bmidat$Program),]

z1 = bmidat[bmidat$Program=="CABIN",]
z1 = z1[order(z1$Perc_of_sites,decreasing=T),]
z1.error = ceiling(max(z1$pct_misclassified,na.rm=T))
z1colours = colorRampPalette(c("grey100","grey1"))(z1.error)

z2 = bmidat[bmidat$Program=="RIVPACS",]
z2 = z2[order(z2$Perc_of_sites,decreasing=T),]
z2.error = ceiling(max(z2$pct_misclassified,na.rm=T))
z2colours = colorRampPalette(c("grey100","grey1"))(z2.error)

z3 = bmidat[bmidat$Program=="AUSRIVAS",]
z3 = z3[order(z3$Perc_of_sites,decreasing=T),]
z3.error = ceiling(max(z3$pct_misclassified,na.rm=T))
z3colours = colorRampPalette(c("grey100","grey1"))(z3.error)


# Label function
label <- function(px, py, lab, ..., adj=c(0, 1)) {
  usr <- par("usr")
  text(usr[1] + px*(usr[2] - usr[1]),
       usr[3] + py*(usr[4] - usr[3]),
       lab, adj=adj, ...)
}

par(mfrow=c(3,1), oma=c(2,2,0.5,0.1),mar=c(2,2,0,0), las=1)
barplot(z1$Perc_of_sites, col=z1colours[z1$pct_misclassified], xaxs="i", ylim=c(0,100), space=0)
legend("right",col="black",
       pt.bg=c("white",z1colours[seq(0,length(z1colours),round(length(z1colours)/8))]),
       pch=22, cex=1.2, pt.cex=2, bty="n",
       legend=seq(0,length(z1colours),round(length(z1colours)/8)) )
label(0.9, 0.5, lab="% misidentified", srt=90, cex=1.5, adj=c(0.5,0.5))
label(0.5, 0.95, lab="CABIN", cex=1.5, adj=c(0.5,0.5))

barplot(z2$Perc_of_sites, col=z2colours[z2$pct_misclassified], xaxs="i", ylim=c(0,100), space=0)
legend("right",col="black",
       pt.bg=c("white",z2colours[seq(0,length(z2colours),round(length(z2colours)/8))]),
       pch=22, cex=1.2, pt.cex=2, bty="n",
       legend=seq(0,length(z2colours),round(length(z2colours)/8)))
label(0.9, 0.5, lab="% misidentified", srt=90, cex=1.5, adj=c(0.5,0.5))
label(0.5, 0.95, lab="RIVPACS", cex=1.5, adj=c(0.5,0.5))
```

```r
barplot(z3$Perc_of_sites, col=z3colours[z3$pct_misclassified], xaxs="i", ylim=c(0,100), space=0)
legend("right",col="black",
       pt.bg=c("white",z3colours[seq(0,length(z3colours),round(length(z3colours)/8))]),
       pch=22, cex=1.2, pt.cex=2, bty="n",
       legend=seq(0,length(z3colours),round(length(z3colours)/8)))
label(0.9, 0.5, lab="% misidentified", srt=90, cex=1.5, adj=c(0.5,0.5))
label(0.5, 0.95, lab="AUSRIVAS", cex=1.5, adj=c(0.5,0.5))

mtext("Frequecy of observation - %", side=2, line=0.5, outer=T, las=0)
```

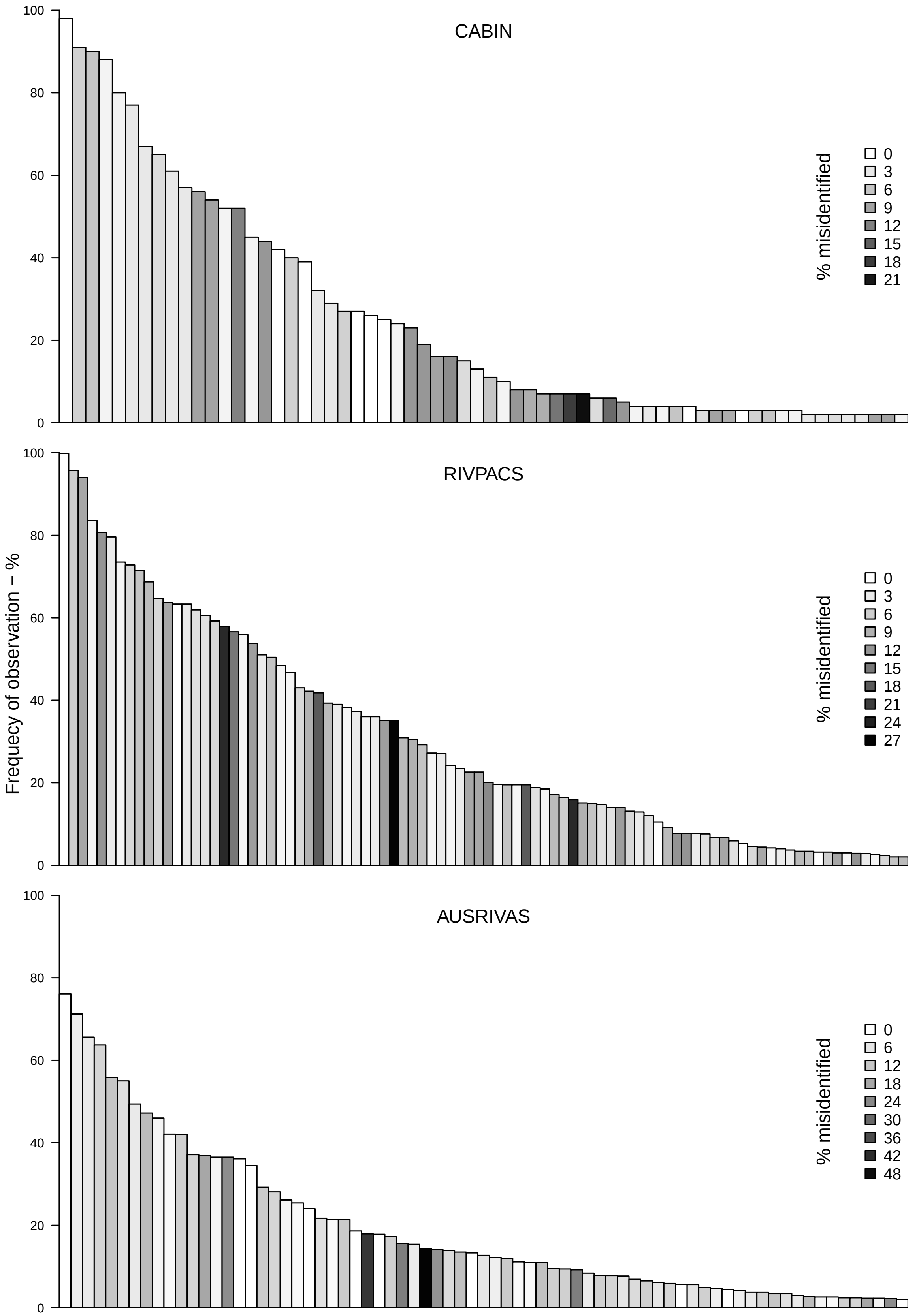
CABIN
RIVPACS
AUSRIVAS
Frequecy of observation – %
% misidentified
0
3
6
9
12
15
18
21
24
27
0
6
12
18
24
30
36
42
48
0
20
40
60
80
100

Data for the CABIN plot:

```
kable(z1, booktabs=TRUE)
```

| | Family | Perc_of_sites | Order | pct_misclassified | Program |
|---|---|---|---|---|---|
| 26 | Chironomidae | 98 | Diptera | 1.3846448 | CABIN |
| 14 | Baetidae | 91 | Ephemeroptera | 5.9076923 | CABIN |
| 69 | Heptageniidae | 90 | Ephemeroptera | 6.8597561 | CABIN |
| 53 | Ephemerellidae | 88 | Ephemeroptera | 2.4475524 | CABIN |
| 27 | Chloroperlidae | 80 | Plecoptera | 2.3809524 | CABIN |
| 102 | Nemouridae | 77 | Plecoptera | 3.8186158 | CABIN |
| 146 | Tipulidae | 67 | Diptera | 3.2325339 | CABIN |
| 111 | Perlodidae | 65 | Plecoptera | 4.2307692 | CABIN |
| 128 | Rhyacophilidae | 61 | Trichoptera | 3.2092426 | CABIN |
| 77 | Hydropsychidae | 57 | Trichoptera | 3.1847134 | CABIN |
| 50 | Empididae | 56 | Diptera | 0.3590664 | CABIN |
| 89 | Leptophlebiidae | 54 | Ephemeroptera | 9.1334895 | CABIN |
| 22 | Capniidae | 52 | Plecoptera | 9.5238095 | CABIN |
| 143 | Taeniopterygidae | 52 | Plecoptera | 2.7777778 | CABIN |
| 135 | Simuliidae | 45 | Diptera | 12.2008049 | CABIN |
| 24 | Ceratopogonidae | 44 | Diptera | 2.0026264 | CABIN |
| 3 | Ameletidae | 42 | Ephemeroptera | 0.0000000 | CABIN |
| 49 | Elmidae | 40 | Coleoptera | 10.3225806 | CABIN |
| 61 | Glossosomatidae | 39 | Trichoptera | 1.5909091 | CABIN |
| 90 | Leuctridae | 32 | Plecoptera | 5.2486188 | CABIN |
| 138 | Sperchontidae | 29 | Trombidiformes | 0.0000000 | CABIN |
| 18 | Brachycentridae | 27 | Trichoptera | 1.5172414 | CABIN |
| 124 | Psychodidae | 27 | Diptera | 3.9071567 | CABIN |
| 85 | Lebertiidae | 26 | Trombidiformes | 0.0000000 | CABIN |
| 86 | Lepidostomatidae | 25 | Trichoptera | 3.0625832 | CABIN |
| 92 | Limnephilidae | 24 | Trichoptera | 5.2262364 | CABIN |
| 110 | Perlidae | 23 | Plecoptera | 1.8813314 | CABIN |
| 147 | Torrenticolidae | 19 | Trombidiformes | 1.3157895 | CABIN |
| 78 | Hydroptilidae | 16 | Trichoptera | 1.5873016 | CABIN |
| 95 | Lumbriculidae | 16 | Lumbriculida | 2.1739130 | CABIN |
| 81 | Hygrobatidae | 15 | Trombidiformes | 10.0000000 | CABIN |
| 116 | Sphaeriidae | 13 | Veneroida | 10.1449275 | CABIN |
| 51 | Enchytraeidae | 11 | Haplotaxida | 9.1575092 | CABIN |
| 149 | Uenoidae | 10 | Trichoptera | 0.4724409 | CABIN |
| 117 | Planariidae | 8 | Tricladida | 0.0000000 | CABIN |
| 126 | Pteronarcyidae | 8 | Plecoptera | 0.7194245 | CABIN |
| 8 | Apataniidae | 7 | Trichoptera | 11.2359551 | CABIN |
| 87 | Leptoceridae | 7 | Trichoptera | 4.5667447 | CABIN |
| 93 | Limnesiidae | 7 | Trombidiformes | 0.0000000 | CABIN |
| 112 | Philopotamidae | 7 | Trichoptera | 2.8023599 | CABIN |
| 118 | Planorbidae | 6 | undef_Gastropoda | 6.6157761 | CABIN |
| 122 | Polycentropodidae | 6 | Trichoptera | 2.3809524 | CABIN |
| 12 | Athericidae | 5 | Diptera | 10.0000000 | CABIN |
| 29 | Coenagrionidae | 4 | Odonata | 8.1812035 | CABIN |
| 58 | Gammaridae | 4 | Amphipoda | 8.0980066 | CABIN |
| 63 | Gomphidae | 4 | Odonata | 13.1799163 | CABIN |
| 114 | Physidae | 4 | undef_Gastropoda | 18.3333333 | CABIN |
| 123 | Psephenidae | 4 | Coleoptera | 0.3649635 | CABIN |
| 6 | Ancylidae | 3 | undef_Gastropoda | 22.7272727 | CABIN |

| | Family | Perc_of_sites | Order | pct_misclassified | Program |
|---|---|---|---|---|---|
| 19 | Caenidae | 3 | Ephemeroptera | 4.8309179 | CABIN |
| 43 | Dixidae | 3 | Diptera | 0.0000000 | CABIN |
| 48 | Dytiscidae | 3 | Coleoptera | 14.8269202 | CABIN |
| 70 | Hyalellidae | 3 | Amphipoda | 0.3246753 | CABIN |
| 74 | Hydrobiidae | 3 | Littorinimorpha | 10.8586831 | CABIN |
| 96 | Lymnaeidae | 3 | undef_Gastropoda | 2.9164330 | CABIN |
| 109 | Peltoperlidae | 3 | Plecoptera | 0.0000000 | CABIN |
| 9 | Arrenuridae | 2 | Trombidiformes | 3.0000000 | CABIN |
| 16 | Blephariceridae | 2 | Diptera | 0.0000000 | CABIN |
| 28 | Chydoridae | 2 | Diplostraca | 2.5125628 | CABIN |
| 32 | Corixidae | 2 | Hemiptera | 6.4171123 | CABIN |
| 33 | Corydalidae | 2 | Megaloptera | 1.8442623 | CABIN |
| 35 | Crangonyctidae | 2 | Amphipoda | 4.0000000 | CABIN |
| 73 | Hydridae | 2 | Anthoathecata | 9.7014925 | CABIN |
| 152 | Valvatidae | 2 | undef_Gastropoda | 8.7719298 | CABIN |

Data for the RIVPACS plot:

```
kable(z2, booktabs=TRUE)
```

| | Family | Perc_of_sites | Order | pct_misclassified | Program |
|---|---|---|---|---|---|
| 127785 | Chironomidae | 99.8 | Diptera | 1 | RIVPACS |
| 79838 | Baetidae | 95.7 | Ephemeroptera | 6 | RIVPACS |
| 60963 | Elmidae | 94.0 | Coleoptera | 10 | RIVPACS |
| 79510 | Ephemerellidae | 83.6 | Ephemeroptera | 2 | RIVPACS |
| 131897 | Simuliidae | 80.7 | Diptera | 12 | RIVPACS |
| 68410 | Hydropsychidae | 79.6 | Trichoptera | 3 | RIVPACS |
| 43683 | Lumbriculidae | 73.5 | Lumbriculida | 2 | RIVPACS |
| 115066 | Leuctridae | 72.8 | Plecoptera | 5 | RIVPACS |
| 79678 | Heptageniidae | 71.5 | Ephemeroptera | 7 | RIVPACS |
| 148574 | Gammaridae | 68.7 | Amphipoda | 8 | RIVPACS |
| 68928 | Limnephilidae | 64.7 | Trichoptera | 5 | RIVPACS |
| 155515 | Sphaeriidae | 63.7 | Veneroida | 10 | RIVPACS |
| 131062 | Pediciidae | 63.3 | Diptera | 1 | RIVPACS |
| 70348 | Rhyacophilidae | 63.3 | Trichoptera | 3 | RIVPACS |
| 114987 | Nemouridae | 61.9 | Plecoptera | 4 | RIVPACS |
| 115341 | Perlodidae | 60.6 | Plecoptera | 4 | RIVPACS |
| 79585 | Caenidae | 59.2 | Ephemeroptera | 5 | RIVPACS |
| 161388 | Ancylidae | 57.9 | undef_Gastropoda | 23 | RIVPACS |
| 49222 | Dytiscidae | 56.6 | Coleoptera | 15 | RIVPACS |
| 69939 | Polycentropodidae | 55.9 | Trichoptera | 2 | RIVPACS |
| 157428 | Hydrobiidae | 53.8 | undef_Gastropoda | 11 | RIVPACS |
| 158452 | Lymnaeidae | 51.0 | undef_Gastropoda | 3 | RIVPACS |
| 43763 | Glossiphoniidae | 50.4 | Arhynchobdellida | 7 | RIVPACS |
| 127227 | Empididae | 48.4 | Diptera | 0 | RIVPACS |
| 115389 | Chloroperlidae | 46.7 | Plecoptera | 2 | RIVPACS |
| 67702 | Glossosomatidae | 43.0 | Trichoptera | 2 | RIVPACS |
| 67963 | Leptoceridae | 42.2 | Trichoptera | 5 | RIVPACS |
| 44567 | Enchytraeidae | 41.8 | Haplotaxida | 9 | RIVPACS |
| 70639 | Sericostomatidae | 39.3 | Trichoptera | 18 | RIVPACS |
| 43890 | Erpobdellidae | 39.0 | Arhynchobdellida | 8 | RIVPACS |
| 70527 | Lepidostomatidae | 38.3 | Trichoptera | 3 | RIVPACS |

| | Family | Perc_of_sites | Order | pct_misclassified | Program |
|---|---|---|---|---|---|
| 69436 | Hydroptilidae | 37.3 | Trichoptera | 2 | RIVPACS |
| 149053 | Asellidae | 36.0 | Isopoda | 3 | RIVPACS |
| 127379 | Limoniidae | 36.0 | Diptera | 2 | RIVPACS |
| 41321 | Planariidae | 35.1 | Tricladida | 0 | RIVPACS |
| 131078 | Tipulidae | 35.1 | Diptera | 3 | RIVPACS |
| 50651 | Haliplidae | 30.9 | Coleoptera | 11 | RIVPACS |
| 51342 | Hydraenidae | 30.5 | Coleoptera | 27 | RIVPACS |
| 61341 | Gyrinidae | 29.2 | Coleoptera | 8 | RIVPACS |
| 79989 | Leptophlebiidae | 27.2 | Ephemeroptera | 9 | RIVPACS |
| 160552 | Planorbidae | 27.1 | undef_Gastropoda | 7 | RIVPACS |
| 79459 | Ephemeridae | 24.2 | Ephemeroptera | 2 | RIVPACS |
| 115315 | Taeniopterygidae | 23.4 | Plecoptera | 3 | RIVPACS |
| 70280 | Goeridae | 22.6 | Trichoptera | 2 | RIVPACS |
| 128723 | Muscidae | 22.6 | Diptera | 3 | RIVPACS |
| 138523 | Hygrobatidae | 20.1 | Trombidiformes | 10 | RIVPACS |
| 133290 | Athericidae | 19.6 | Diptera | 10 | RIVPACS |
| 65964 | Helophoridae | 19.5 | Coleoptera | 13 | RIVPACS |
| 115265 | Perlidae | 19.5 | Plecoptera | 2 | RIVPACS |
| 71256 | Sialidae | 19.5 | Megaloptera | 7 | RIVPACS |
| 69641 | Psychomyiidae | 18.8 | Trichoptera | 2 | RIVPACS |
| 76882 | Veliidae | 18.5 | Hemiptera | 0 | RIVPACS |
| 160044 | Physidae | 17.1 | undef_Gastropoda | 18 | RIVPACS |
| 132224 | Psychodidae | 16.4 | Diptera | 4 | RIVPACS |
| 43837 | Piscicolidae | 15.9 | Arhynchobdellida | 0 | RIVPACS |
| 70485 | Brachycentridae | 15.1 | Trichoptera | 2 | RIVPACS |
| 44429 | Lumbricidae | 15.0 | Haplotaxida | 8 | RIVPACS |
| 138488 | Lebertiidae | 14.7 | Trombidiformes | 0 | RIVPACS |
| 73124 | Corixidae | 14.0 | Hemiptera | 6 | RIVPACS |
| 51661 | Scirtidae | 14.0 | Coleoptera | 23 | RIVPACS |
| 162726 | Valvatidae | 13.1 | undef_Gastropoda | 9 | RIVPACS |
| 163053 | Bithyniidae | 12.9 | undef_Gastropoda | 7 | RIVPACS |
| 147680 | Crangonyctidae | 12.0 | Amphipoda | 4 | RIVPACS |
| 138911 | Sperchontidae | 10.5 | Trombidiformes | 0 | RIVPACS |
| 114095 | Calopterygidae | 9.2 | Odonata | 4 | RIVPACS |
| 129745 | Dixidae | 7.7 | Diptera | 0 | RIVPACS |
| 70098 | Odontoceridae | 7.7 | Trichoptera | 11 | RIVPACS |
| 129447 | Tabanidae | 7.7 | Diptera | 5 | RIVPACS |
| 70681 | Philopotamidae | 7.6 | Trichoptera | 3 | RIVPACS |
| 157164 | Neritidae | 6.8 | Cycloneritida | 4 | RIVPACS |
| 41302 | Dugesiidae | 6.7 | Tricladida | 1 | RIVPACS |
| 113709 | Coenagrionidae | 5.9 | Odonata | 8 | RIVPACS |
| 74785 | Aphelocheiridae | 5.2 | Hemiptera | 0 | RIVPACS |
| 75735 | Gerridae | 4.6 | Hemiptera | 12 | RIVPACS |
| 48920 | Hydrophilidae | 4.4 | Coleoptera | 12 | RIVPACS |
| 114956 | Cordulegastridae | 4.2 | Odonata | 3 | RIVPACS |
| 74745 | Notonectidae | 4.0 | Hemiptera | 4 | RIVPACS |
| 161634 | Acroloxidae | 3.7 | undef_Gastropoda | 5 | RIVPACS |
| 69425 | Beraeidae | 3.4 | Trichoptera | 0 | RIVPACS |
| 115111 | Capniidae | 3.4 | Plecoptera | 10 | RIVPACS |
| 130691 | Ptychopteridae | 3.2 | Diptera | 4 | RIVPACS |
| 130714 | Stratiomyidae | 3.2 | Diptera | 1 | RIVPACS |
| 67912 | Molannidae | 3.0 | Trichoptera | 5 | RIVPACS |

| | Family | Perc_of_sites | Order | pct_misclassified | Program |
|---|---|---|---|---|---|
| 154928 | Unionidae | 3.0 | Unionoida | 10 | RIVPACS |
| 79657 | Siphlonuridae | 2.9 | Ephemeroptera | 2 | RIVPACS |
| 70168 | Phryganeidae | 2.8 | Trichoptera | 3 | RIVPACS |
| 104329 | Crambidae | 2.6 | Lepidoptera | 3 | RIVPACS |
| 77107 | Micronectidae | 2.4 | Hemiptera | 0 | RIVPACS |
| 164822 | Gastrodontidae | 2.0 | Stylommatophora | 8 | RIVPACS |
| 163595 | Succineidae | 2.0 | Stylommatophora | 7 | RIVPACS |

Data for the AUSRIVAS plot:

```
kable(z3, booktabs=TRUE)
```

| | Family | Perc_of_sites | Order | pct_misclassified | Program |
|---|---|---|---|---|---|
| 17 | Chironomidae | 76.1 | Diptera | 1.3846448 | AUSRIVAS |
| 671 | Leptoceridae | 71.2 | Trichoptera | 4.5667447 | AUSRIVAS |
| 910 | Baetidae | 65.6 | Ephemeroptera | 5.9076923 | AUSRIVAS |
| 68 | Leptophlebiidae | 63.7 | Ephemeroptera | 9.1334895 | AUSRIVAS |
| 107 | Simuliidae | 55.8 | Diptera | 12.2008049 | AUSRIVAS |
| 46 | Gripopterygidae | 55.0 | Plecoptera_Insecta | 7.9925651 | AUSRIVAS |
| 551 | Hydrobiosidae | 49.4 | Trichoptera | 5.5363322 | AUSRIVAS |
| 321 | Dytiscidae | 47.2 | Coleoptera | 14.8269202 | AUSRIVAS |
| 59 | Hydropsychidae | 46.0 | Trichoptera | 3.1847134 | AUSRIVAS |
| 7 | Atyidae | 42.1 | Decapoda | 2.6915114 | AUSRIVAS |
| 341 | Elmidae | 42.0 | Coleoptera | 10.3225806 | AUSRIVAS |
| 1 | Aeshnidae | 37.1 | Odonata | 9.9216710 | AUSRIVAS |
| 121 | Veliidae | 36.9 | Hemiptera | 0.0000000 | AUSRIVAS |
| 931 | Physidae | 36.5 | undef_Gastropoda | 18.3333333 | AUSRIVAS |
| 120 | Tipulidae | 36.5 | Diptera | 3.2325339 | AUSRIVAS |
| 104 | Scirtidae | 36.1 | Coleoptera | 23.0312036 | AUSRIVAS |
| 241 | Corydalidae | 34.5 | Megaloptera | 1.8442623 | AUSRIVAS |
| 161 | Chiltoniidae | 29.2 | Amphipoda | 1.0791367 | AUSRIVAS |
| 1001 | Psephenidae | 28.1 | Coleoptera | 0.3649635 | AUSRIVAS |
| 581 | Hydrophilidae | 26.1 | Coleoptera | 11.7822521 | AUSRIVAS |
| 201 | Conoesucidae | 25.4 | Trichoptera | 9.6000000 | AUSRIVAS |
| 80 | Notonectidae | 24.0 | Hemiptera | 3.6764706 | AUSRIVAS |
| 15 | Ceratopogonidae | 21.7 | Diptera | 2.0026264 | AUSRIVAS |
| 441 | Glossosomatidae | 21.4 | Trichoptera | 1.5909091 | AUSRIVAS |
| 471 | Gyrinidae | 21.4 | Coleoptera | 7.6923077 | AUSRIVAS |
| 901 | Philopotamidae | 18.6 | Trichoptera | 2.8023599 | AUSRIVAS |
| 291 | Dixidae | 17.9 | Diptera | 0.0000000 | AUSRIVAS |
| 41 | Gerridae | 17.8 | Hemiptera | 11.8387909 | AUSRIVAS |
| 601 | Hydroptilidae | 17.2 | Trichoptera | 1.5873016 | AUSRIVAS |
| 810 | Austroperlidae | 15.6 | Plecoptera | 40.0000000 | AUSRIVAS |
| 811 | Notonemouridae | 15.4 | Plecoptera | 1.8181818 | AUSRIVAS |
| 381 | Eustheniidae | 14.3 | Plecoptera | 0.0000000 | AUSRIVAS |
| 911 | Philorheithridae | 14.1 | Trichoptera | 10.4477612 | AUSRIVAS |
| 1021 | Ptilodactylidae | 13.9 | Coleoptera | 26.9230769 | AUSRIVAS |
| 331 | Ecnomidae | 13.5 | Trichoptera | 5.4878049 | AUSRIVAS |
| 1131 | Synlestidae | 13.3 | Odonata | 50.0000000 | AUSRIVAS |
| 310 | Ancylidae | 12.7 | undef_Gastropoda | 22.7272727 | AUSRIVAS |
| 181 | Coenagrionidae | 12.2 | Odonata | 8.1812035 | AUSRIVAS |
| 141 | Calocidae | 12.0 | Trichoptera | 0.0000000 | AUSRIVAS |

| | Family | Perc_of_sites | Order | pct_misclassified | Program |
|---|---|---|---|---|---|
| 45 | Gomphidae | 11.1 | Odonata | 13.1799163 | AUSRIVAS |
| 351 | Empididae | 10.9 | Diptera | 0.3590664 | AUSRIVAS |
| 741 | Micronectidae | 10.9 | Hemiptera | 0.0000000 | AUSRIVAS |
| 64 | Atriplectididae | 9.5 | Trichoptera | 0.0000000 | AUSRIVAS |
| 511 | Helicopsychidae | 9.4 | Trichoptera | 1.9867550 | AUSRIVAS |
| 221 | Corixidae | 9.2 | Hemiptera | 6.4171123 | AUSRIVAS |
| 125 | Caenidae | 8.4 | Ephemeroptera | 4.8309179 | AUSRIVAS |
| 271 | Cyrenidae | 7.9 | Veneroida | 0.0000000 | AUSRIVAS |
| 83 | Odontoceridae | 7.8 | Trichoptera | 11.3793103 | AUSRIVAS |
| 98 | Pontogeneiidae | 7.7 | Amphipoda | 2.0000000 | AUSRIVAS |
| 961 | Polycentropodidae | 6.9 | Trichoptera | 2.3809524 | AUSRIVAS |
| 56 | Hydrochidae | 6.5 | Coleoptera | 13.4831461 | AUSRIVAS |
| 501 | Helicophidae | 6.1 | Trichoptera | 10.8695652 | AUSRIVAS |
| 881 | Paramelitidae | 5.9 | Amphipoda | 0.0000000 | AUSRIVAS |
| 541 | Hydrobiidae | 5.7 | undef_Gastropoda | 10.8586831 | AUSRIVAS |
| 52 | Hydraenidae | 5.6 | Coleoptera | 26.9826800 | AUSRIVAS |
| 103 | Pyralidae | 4.9 | Lepidoptera | 5.4794521 | AUSRIVAS |
| 5 | Athericidae | 4.7 | Diptera | 10.0000000 | AUSRIVAS |
| 691 | Lestidae | 4.4 | Odonata | 9.3750000 | AUSRIVAS |
| 94 | Planorbidae | 4.2 | undef_Gastropoda | 6.6157761 | AUSRIVAS |
| 211 | Corduliidae | 3.8 | Odonata | 8.3333333 | AUSRIVAS |
| 1081 | Sphaeriidae | 3.8 | Veneroida | 10.1449275 | AUSRIVAS |
| 31 | Dugesiidae | 3.4 | Tricladida | 0.5050505 | AUSRIVAS |
| 65 | Janiridae | 3.4 | Isopoda | 6.2500000 | AUSRIVAS |
| 191 | Coloburiscidae | 3.0 | Ephemeroptera | 0.0000000 | AUSRIVAS |
| 871 | Paracalliopiidae | 2.7 | Amphipoda | 0.0000000 | AUSRIVAS |
| 531 | Hydridae | 2.6 | Anthoathecata | 9.7014925 | AUSRIVAS |
| 62 | Hymenosomatidae | 2.6 | Decapoda | 0.0000000 | AUSRIVAS |
| 951 | Pleidae | 2.4 | Hemiptera | 0.0000000 | AUSRIVAS |
| 1121 | Stratiomyidae | 2.4 | Diptera | 1.2048193 | AUSRIVAS |
| 841 | Oniscigastridae | 2.3 | Ephemeroptera | 0.0000000 | AUSRIVAS |
| 861 | Palaemonidae | 2.3 | Decapoda | 6.8466731 | AUSRIVAS |
| 261 | Culicidae | 2.2 | Diptera | 10.4496951 | AUSRIVAS |
| 761 | Naucoridae | 2.0 | Hemiptera | 4.8780488 | AUSRIVAS |

```r
### >>>  NB  <<<
# Thinking further about how we summarise the cumulative rates of error as a percentage of Families
# I tried to standardize by richness. However, by standardizing by richness, all columns effectively
# become the same so the total rate of misclassification can be communicated by a single value.

cumulative.misclassifications.func = function(data, sims=1000){
  zmat = matrix(NA, nrow=sims, ncol=nrow(data))
  for(i in 1:sims){
    for(j in 1:nrow(data)){
      pj        = 1-(data$pct_misclassified[sample(1:nrow(data), j, prob=data$Perc_of_sites, replace=F)
      zmat[i,j] = sum(rbinom(n=j, size=1, prob=pj))
    }
  }
  # No. of errors
  zmat = matrix(rep(c(1:ncol(zmat)),nrow(zmat)),ncol=ncol(zmat),byrow=T) - zmat
  # Errors as percentage of richness
  zmat = zmat / (matrix(rep(c(1:ncol(zmat)),nrow(zmat)),ncol=ncol(zmat),byrow=T)/100)
```

```r
  zmat = apply(zmat, 2, mean)
  mean(zmat)
}
z1.errormean = cumulative.misclassifications.func(z1, 10000)
z2.errormean = cumulative.misclassifications.func(z2, 10000)
z3.errormean = cumulative.misclassifications.func(z3, 10000)

round(c(z1.errormean,z2.errormean,z3.errormean),2)
```

```
## [1] 4.45 6.11 7.78
```

So for CABIN the overall rate of Family misidentification is currently 4.45%, RIVPACS it is 6.09% and AUSRIVAS it is 7.79%. Clarke 2009 reported an error rate in UK around 8% for morphologically identified Families and although a comparable figure was not as easy to draw fom Haase et al. 2006, the rate was probably as high if not higher. DNA reference databases have only begun to be actively developed in the last 8 years with the establishment of the Barcode of Life project, and its continued expansion to better represent the genetic diversity within and among species is likely to continue to reduce the rates of misclassification using DNA barcoding.
