## Supplement 2 for "Studying ecosystems with DNA metabarcoding: lessons from aquatic biomonitoring"

### Supplementary Material S2: Comparing observations of freshwater macroinvertebrate families based on morphological identification and DNA metabarcoding.

*Alex Bush*

*25th July 2018*

This supplementary file provides a preliminary analysis of benthic macroinvertebrate samples from Atlantic Canada as part of the the Government of Canada's Genomic Research and Development Initiative.

---

#### Morphological Data

As part of the Ecobiomics project parallel samples were collected and processed for metabarcoding as part of the Canadian Aquatic Biomonitoring Network (CABIN). Data were collected following procedures outlined in detail in the CABIN website:
https://www.canada.ca/en/environment-climate-change/services/canadian-aquatic-biomonitoring-network.html

We would like to acknowledge and thank our partners and colleagues in Atlantic Canada who helped collect additional samples for metabarcoding and have allowed us to use the morphological data from their monitoring studies:
> Abegweit Conservation Society
> Bluenose Coastal Action Group
> Cape Breton Atlantic Coastal Action Program
> Central Queens Wildlife Federation, PEI
> Department of National Defence at Gagetown
> Fort Folly Habitat Recovery Project
> Miramichi River Environmental Assessment Committee
> Morell River Management Cooperative
> Parks Canada (Fundy, Kouchibouguac, Cape Breton, Kejimkujik and Forillon National Parks)
> PEI department of Communities, Land and Environment
> Petitcodiac Watershed Alliance
> Souris and Area Branch of the PEI Wildlife Federation
> Southern Gulf of St.Lawrence Coalition
> Tabusintac Watershed Association
> Unama'ki Institute of Natural Resources for the Bras d'Or region

Finally many thanks to staff at Environment and Climate Change Canada, Natural Resources Canada and the Department for Fisheries and Oceans.

#### DNA metabarcoded data

##### Collection of bulk-community samples

Samples were collected for metabarcoding using the same standard protocols as the CABIN morphological samples. The primary modifications to traditional protocols include treating sampling equipment with diluted household bleach (1 in 10 dilution resulting in about 0.5% sodium hypochlorite) between surveys and during sample processing to avoid cross-contamination. After collection, samples destined for sequencing should be stored in 95% ethanol and then as a precaution, kept as cold as possible in the field, during storage and in shipping to prevent the DNA from degrading.

##### DNA extraction, amplification and sequencing

After removing rocks, macroinvertebrate samples were homogenized using a WaringPro® blender. DNA was extracted using the DNeasy PowerSoil Kit (Qiagen, Catalog No. 12888-100) following the manufacturers instructions (available online).

Three loci from the COI barcode region were amplified. The primers used in the current workflow include the F230 primers from *Gibson et al. (2015, PLoS ONE, 10, e0138432)*, although the reverse primer was modified to use "N" rather than inosine (I) because the PCR product for the former performed better on agarose gel. We also used fwh2 and BF2-BR2 primers (*Elbrecht & Leese 2017 Frontiers in Environmental Science, 5; Vamos, Elbrecht & Leese 2017 Metabarcoding and Metagenomics, 1*), because they each identified a significant unique proportion of the community that was complementary to the other, and to F230 (i.e. a number of taxa were only identified by one primer pair).

| Amplicon | Primer_name | Forward_5_.._3 |
|---|---|---|
| BF2-BR2 | BF2 | GCHCCHGAYATRGCHTTYCC |
| BF2-BR2 | BR2 | TCDGGRTGNCCRAARAAYCA |
| Fwh2 | Fwh2 F. | GGDACWGGWTGAACWGTWTAYCCHCC |
| Fwh2 | Fwh2 R. | GTRATWGCHCCDGCTARWACWGG |
| F230 | LCO1490-Folmar-F | GGTCAACAAATCATAAAGATATTGG |
| F230 | F230R_modN | CTTATRTTRTTTATNCGNGGRAANGC |

A two-step PCR approach was used to prepare the Illumina-competent amplicon libraries. During the first PCR, each sample was amplified for F230, fwh2 and BF2-BR2 in separate reactions (giving approximate fragment sizes of 347 bp, 321 bp and 528 bp respectively) using Amplitaq Gold 360 Mastermix (Applied Biosystems, Catalog No. 4398881) in a total volume of 25 µl. A total of 5 (fwh2) or 6 (BF-BR2 and F230) oligonucleotides consisting of the locus-specific and Illumina-tailed locus-specific with or without off-setting bases to increase diversity were used in each separate reaction for the three loci. F230 and fwh2 were amplified following these PCR conditions: initial incubation at 95°C for 10 min; 30 cycles of denaturation at 95°C for 30 sec; annealing at 46°C for 30 sec; extension at 60°C for 60 sec; and, a final extension at 72°C for 7 min. BF2-BR2 was amplified using the same conditions except the annealing was at 48°C instead of 46°C.

The second round of PCR used the Q5 Hot start High-fidelity DNA polymerase (New England BioLabs, Catalog No.M0493S) for all three amplicons. Unique forward and reverse combinations of Nextera XT dual 8 bp index sequences were used for each sample, which enables bioinformatic de-multiplexing of individual samples. Reactions were performed using 12.0 uL of H2O, 0.25 µL of Q5 Hot start High-fidelity DNA polymerase, 5 µL of 5X Q5 reaction buffer, 0.5 µL of 10 nM dNTPs, 0.625 µL of Forward Index primer (20 µm stock concentration), 0.625 µL of Reverse Index primer (20 µm stock concentration), 1 uL of 1% BSA and 5 uL of SPRI bead-cleaned DNA from PCR step 1. Each amplicon was amplified separately under the following conditions: initial incubation at 98°C for 30 sec; 10 cycles of denaturation at 98°C for 15 sec; annealing at 66°C for 20 sec; extension at 72°C for 30 sec; and, a final extension at 72°C for 2 min.

Samples were normalized using the SequalPrep Normalization Plate (ThermoFisher Scientific, Catalog. No. A1051001) following the manufacturer's protocol. Pooled samples were cleaned using a ratio of 1.2x of High Prep PCR magnetic beads following the manufacturer's protocol as mentioned above. DNA was resuspended in 40 uL of Tris-HCl 10 mM, pH 8.0. Finally samples were sequenced on using a MiSeq V2 2 x 250 bp run with a 15% PhiX spike-in to increase base diversity.

##### Bioinformatics

Sequencing reads were analyzed using in-house pipelines and are available online: https://github.com/EcoBiomics-Zoobiome/SCVUC_COI_metabarcode_pipeline The **SCVUC pipeline** refers to the programs, algorithms, and reference datasets used in this data flow: SEQPREP, CUTADAPT, VSEARCH, UNOISE and the COI classifier (Porter & Hajibabaei, 2018 Sci Rep).

---

#### Consistency between paired kick samples

The supplementary file contains number of detections of each taxon using morphological and DNA metabarcoding approaches, and an estimate of the detectability of each taxon in a kick samples based on the similarity of replicate kick samples (see Figure 1a main text).

```
fx = read.csv("Supplementary_file_3_Morphological_and_DNA_macroinvertebrate_data_for_Figure_4.csv")
```

First we can reproduce Figure 4 in the main text. The top row (a and b) indicates the correspondance between the detection of a given macroinveretbrate family using morphological identification, and its detection in **a seperate paired kick-sample** using metabarcoding. Larger circles indicate taxa observed most often, and we can see that typically the majority of taxa observed by morphology are also observed at the same sites using metabarcoding.

Points that fall below the diagonal indicate taxa that were detected more readily by DNA than by morphology - specifically the proportion of detections made by morphology that were matched by DNA exceeded the proportion of detections made by DNA matched by morphology.

```
A         = 5 # Taxon observed by Morphology only column
B         = 6 # Taxon observed by DNA only column
AB        = 7 # Taxon observed by both Morphology and DNA column
X         = 8 # Neither Morphology or DNA observed taxon column
cap.prob  = 9 # Probability of occurrence within replicate samples

par(mfrow=c(1,2), las=1,mar=c(4.1,4.1,0.1,0.5), mgp=c(2.2,1,0))

# Explanatory panel I
plot(c(0,100),c(0,100), col="white",
     xlab = "Morphology matched by DNA - %", ylab = "DNA matched by morphology - %")
abline(0,1,lty=3)
points(c(5,5,95,95),c(5,95,5,95), pch=21, cex=5, bg="grey66")
text(60,15, adj=c(0,0), cex=0.9,
     labels="DNA consistently detects \nmorphological records, but \nmorphology detected few \nof the D
text(0,75,  adj=c(0,0), cex=0.9,
     labels="Morphology consistently \ndetects DNA records, \nbut DNA detected few of \nthe morphology
text(55,85, adj=c(0,0), cex=0.9,
     labels="Taxa are consistently \nobserved by both \nmorphological and \nDNA approaches.")
text(18,0, adj=c(0,0), cex=0.9,
     labels="Taxa are rarely \nobserved by both \nmorphological and \nDNA approaches.")
mtext("a)",2,2.5,outer=F,at=100,las=1,cex=1.3)

# Matching Morphological and DNA observations
morph.vs.dna.plot(xdat = fx, A, B, AB, X)
mtext("b)",2,2.5,outer=F,at=100,las=1,cex=1.3)
```

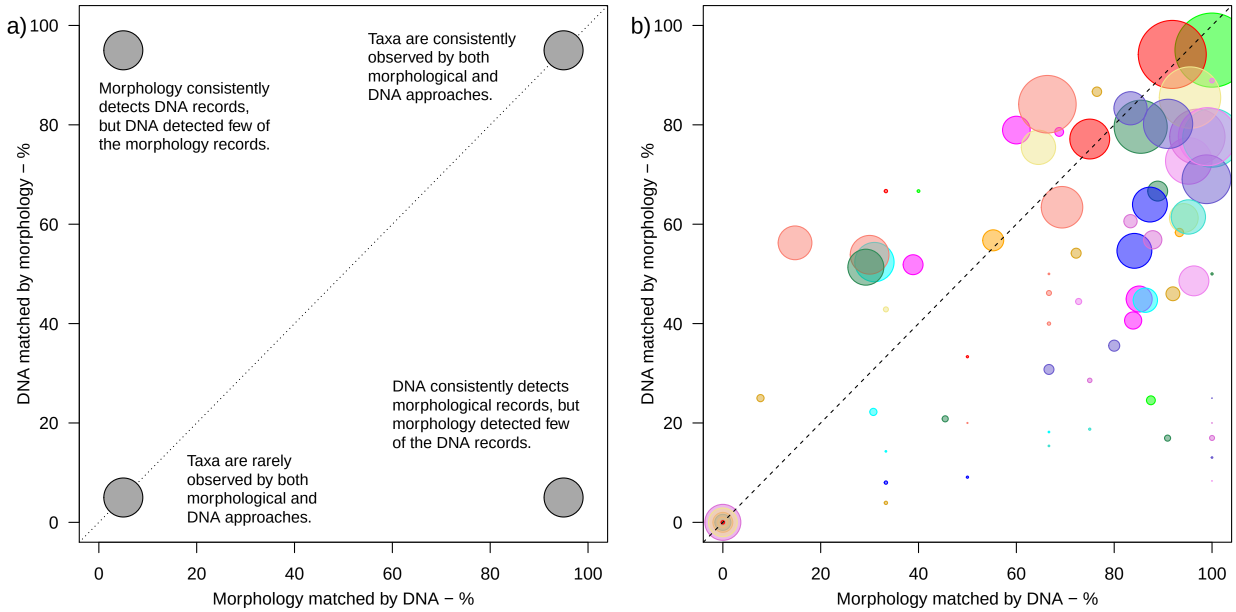


Below are the taxa that occur in the bottom-left corner of a) and b) panels above because the observations of each method did not overlap. This occurred when taxa were observed with one method but not the other, but also when both methods recorded the taxa, just not at the same sites. Taxa that appear to be consistently overlooked by DNA are likely to be due to gaps in the reference library (aquatic mites and oligochaetes in particular) and primer bias (*Elbrecht et al. 2017 Ecology and Evolution, 7*).

| | Class | Order | Family | Morphs_only_sum | DNA_only_sum | Morph_and_DNA |
|---|---|---|---|---|---|---|
| 7 | Arachnida | Trombidiformes | Aturidae | 36 | 0 | 0 |
| 11 | Insecta | Odonata | Calopterygidae | 2 | 3 | 0 |
| 12 | Malacostraca | Decapoda | Cambaridae | 0 | 3 | 0 |
| 13 | Ostracoda | Podocopida | Candonidae | 0 | 23 | 0 |
| 15 | Insecta | Coleoptera | Carabidae | 0 | 36 | 0 |
| 17 | Insecta | Diptera | Chaoboridae | 0 | 1 | 0 |
| 20 | Insecta | Coleoptera | Chrysomelidae | 0 | 2 | 0 |
| 21 | Branchiopoda | Diplostraca | Chydoridae | 0 | 2 | 0 |
| 23 | Insecta | Odonata | Cordulegastridae | 2 | 15 | 0 |
| 27 | Insecta | Lepidoptera | Crambidae | 0 | 6 | 0 |
| 28 | Malacostraca | Amphipoda | Crangonyctidae | 2 | 0 | 0 |
| 29 | Insecta | Diptera | Culicidae | 0 | 14 | 0 |
| 30 | Insecta | Coleoptera | Curculionidae | 0 | 4 | 0 |
| 31 | Maxillopoda | Cyclopoida | Cyclopidae | 0 | 1 | 0 |
| 32 | Branchiopoda | Diplostraca | Daphniidae | 0 | 1 | 0 |
| 33 | Insecta | Trichoptera | Dipseudopsidae | 0 | 3 | 0 |
| 34 | Insecta | Diptera | Dixidae | 3 | 0 | 0 |
| 35 | Insecta | Diptera | Dolichopodidae | 0 | 1 | 0 |
| 36 | Insecta | Coleoptera | Dytiscidae | 1 | 1 | 0 |
| 39 | Clitellata | NA | Enchytraeidae | 29 | 0 | 0 |
| 43 | Insecta | Diptera | Ephydridae | 2 | 3 | 0 |
| 44 | Arachnida | Trombidiformes | Feltriidae | 16 | 0 | 0 |
| 46 | Insecta | Hemiptera | Gerridae | 0 | 3 | 0 |
| 49 | Insecta | Trichoptera | Goeridae | 3 | 13 | 0 |
| 56 | Arachnida | Trombidiformes | Hydrachnidae | 3 | 0 | 0 |

Because the comparison is being made between the detections of taxa in seperate paired samples, there is the possibility that mismatches are genuine, and that samples differred in the taxa they collected by chance (see main text for full discussion). As such, rather than judging an approach based on whether it is able to replicate the detections of the other method, panel d) provides a guide to the likelihood of a taxon being falsely assigned as absent using a particular approach.

```r
par(mfrow=c(1,2), las=1,mar=c(4.1,4.1,0.1,0.5), mgp=c(2.2,1,0))

# Explanatory panel II
plot(c(0,1),c(0,1), col="white",
     xlab = "p(false absence with DNA)", ylab = "p(false absence with morphology)")
abline(0,1,lty=3)
points(c(0.05,0.05,0.95,0.95),c(0.05,0.95,0.05,0.95), pch=21, cex=5, bg="grey66")
text(0.55,0.80, adj=c(0,0), cex=0.9,
     labels="High likelihood both \nmorphological and \nDNA approaches \nincluded a false \nabsence")
text(0,0.75,  adj=c(0,0), cex=0.9,
     labels="High likelihood morphology, \nbut not DNA data \nincluded a false absence")
text(0.15,0, adj=c(0,0), cex=0.9,
     labels="Low probability of \nfalse absences in either \nmorphological or DNA data")
text(0.6,0.15, adj=c(0,0), cex=0.9,
     labels="High likelihood DNA, \nbut not morphology data\nincluded a false absence")
mtext("c)",2,2.5,outer=F,at=1,las=1,cex=1.3)

# Probability a taxon was missed at least once among the remaining samples
# (i.e. observations include at least 1 false absence)
morph.vs.dna.plot(xdat = fx, A, B, AB, X, cap.prob)
mtext("d)",2,2.5,outer=F,at=1,las=1,cex=1.3)
```

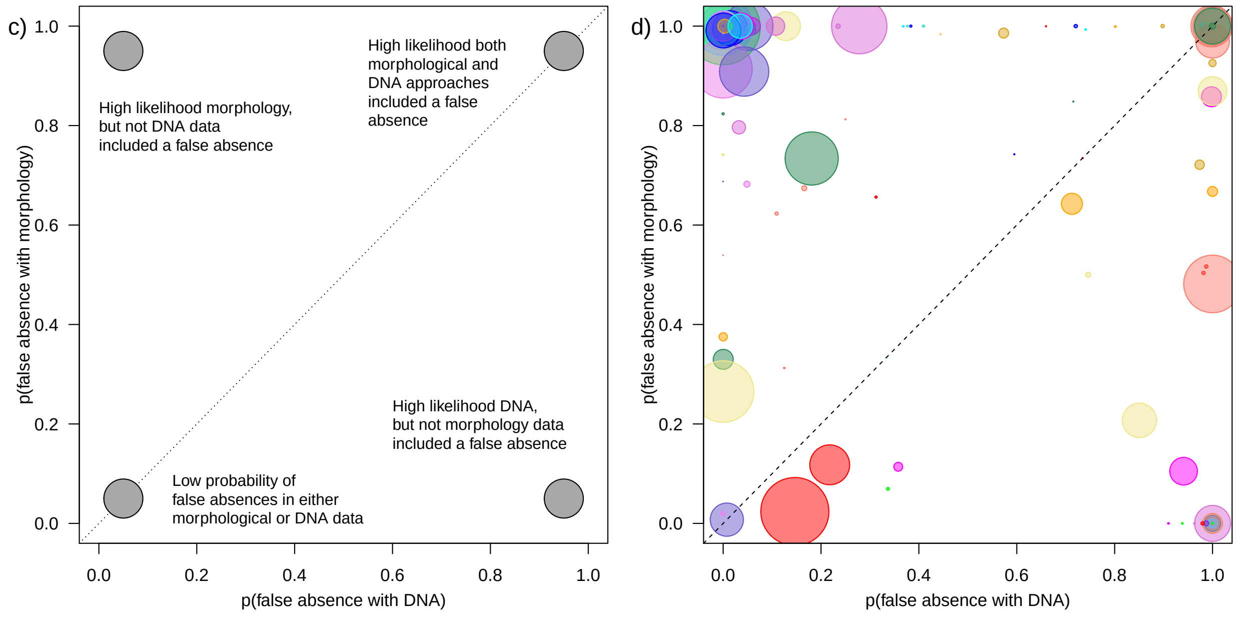


The figure above highlights the likelihood that observations for a particular taxon included **at least one** false absence. However, this is quite a strict threshold for describing error. How are taxa distributed when we consider the likelihood of multiple (1,2,3,4,6,8) false-absences?

```
# Pick some light bright colours to help see overlaps
colours = colors()[grep("dark",colors())]
colours = unique(gsub("dark","",colours[-c(grep('[0-9]',colours),grep("gray",colours),grep("grey",colou
cols    = colours[rep(seq(1,length(colours),1),(nrow(fx)/length(colours))+1)][1:nrow(fx)] #colours[samp

par(mfrow=c(3,2), mar=c(4,4,0.1,0.1),mgp=c(2.3,1,0))
# Compare the probability the Morphological and DNA datasets missed occurrences of a Family
# once, twice, three, four, six and eight times.
for(i in c(1,2,3,4,6,8)){
  Xaxis = 1-pbinom(fx[,AB], size=(ifelse(fx[,A]%in%c(0:i),0,(fx[,A]-i))+fx[,AB]), prob=fx[,cap.prob])
  Xlab  = paste0("p(DNA includes ",i," false absences)") ; Xlim = c(0,1)
  # Yaxis
  Yaxis = 1-pbinom(fx[,AB], size=(ifelse(fx[,B]%in%c(0:i),0,(fx[,B]-i))+fx[,AB]), prob=fx[,cap.prob])
  Ylab  = paste0("p(Morphology includes ",i," false absences)") ; Ylim =c(0,1)
  plot( Yaxis ~ Xaxis, pch=21, col=cols, cex = 1,
        xlim = Xlim, ylim = Ylim, bg = rgb(t(col2rgb(cols)[1:3,]/255),alpha=0.5),
        xlab = Xlab, ylab = Ylab)
  abline(0,1,lty=2)
  points(mean(Xaxis),mean(Yaxis),pch=4,cex=2)
}
```

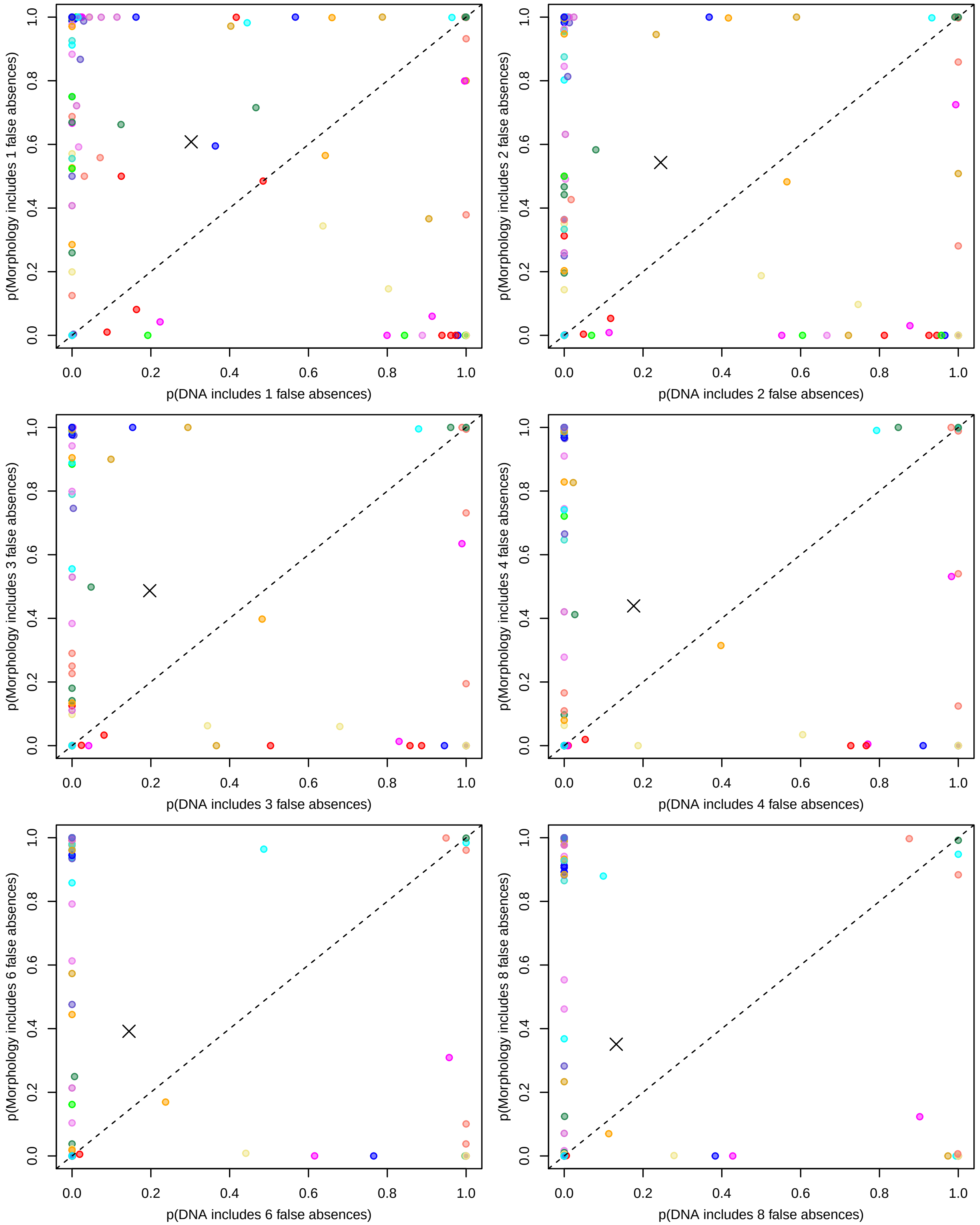


The 'X' on each plot indicates the overall mean, and confirms the sense that false absences for freshwater macroinvertebrates are more likely to be occurring within the morphological dataset.
